# Neotenic expansion of adult-born dentate granule cells reconfigures GABAergic inhibition to enhance social memory consolidation

**DOI:** 10.1101/2025.03.17.643806

**Authors:** Ain Chung, Jason Bondoc Alipio, Megha Ghosh, Liam Evans, Samara M. Miller, Travis D. Goode, Iyanah Mehta, Omar J. Ahmed, Amar Sahay

## Abstract

Adult-born dentate granule cells (abDGCs) contribute to hippocampal dentate gyrus (DG)-CA3/CA2 circuit functions in memory encoding, retrieval and consolidation. Heightened synaptic and structural plasticity of immature abDGCs is thought to govern their distinct contributions to circuit and network mechanisms of hippocampal-dependent memory operations. Protracted maturation or neoteny of abDGCs in higher mammals is hypothesized to offset decline in adult hippocampal neurogenesis by expanding the capacity for circuit and network plasticity underlying different memory operations. Here, we provide evidence for this hypothesis by genetically modelling the effective impact of neoteny of abDGCs on circuitry, network properties and social cognition in mice. We show that selective synchronous expansion of a single cohort of 4 weeks old immature, but not 8 weeks old mature abDGCs, increases functional recruitment of fast spiking parvalbumin expressing inhibitory interneurons (PV INs) in CA3/CA2, number of PV IN-CA3/CA2 synapses, and GABAergic inhibition of CA3/CA2. This transient increase in feed-forward inhibition in DG-CA2 decreased social memory interference and enhanced social memory consolidation. *In vivo* local field potential recordings revealed that the expansion of a single cohort of 4-week-old abDGCs increased baseline power, amplitude, and duration, as well as sensitivity to social investigation-dependent rate changes of sharp-wave ripples (SWRs) in CA1 and CA2, a neural substrate for memory consolidation. Inhibitory neuron-targeted chemogenetic manipulations implicate CA3/CA2 INs, including PV INs, as necessary and sufficient for social memory consolidation following neotenic expansion of the abDGC population and in wild-type mice, respectively. These studies suggest that neoteny of abDGCs may represent an evolutionary adaptation to support cognition by reconfiguring PV IN-CA3/CA2 circuitry and emergent network properties underlying memory consolidation.

## Main

Neurogenesis in the dentate gyrus (DG) subregion of the hippocampus generates dentate granule cells, DGCs, throughout life^1-6^. Adult-born dentate granule cells (abDGCs) functionally integrate into hippocampal circuitry ^7^ and contribute to memory encoding, retrieval and consolidation^8^. Immature abDGCs are characterized by unique electrophysiological properties, connectivity and synaptic plasticity^9-23^ that are thought to underlie their distinct contributions to circuitry ^24-28,29,30^, network properties ^8,31-39^ and memory functions ^16,32,37,39-47^. In mice, the best studied model for adult hippocampal neurogenesis, these defining characteristics of immature abDGC state are most pronounced in 4 weeks old abDGCs and are diminished in 8 weeks old abDGCs ^7,8^. The extent to which hippocampal neurogenesis persists during the lifespan of higher mammals is debated ^6^. Direct and indirect evidence from non-human primates^48,49^ and humans ^50-55^, respectively, suggests that hippocampal neurogenesis drops precipitously in early life but that abDGCs exhibit protracted maturation or neoteny such that abDGCs persist in an immature cell state for an extended period of time. Neoteny of abDGCs in higher mammals may compensate for the decline in adult hippocampal neurogenesis by maintaining a reservoir of highly plastic, experience-modifiable immature abDGCs to support memory operations of the DG-CA3/CA2 circuit in cognition ^53,56^.

Studies in humans ^57-59^, non-human primates^60^ and rodents^61-68^ suggest an important role for the DG in decreasing memory interference^69,70^. Consistently, many studies in rodents have functionally implicated abDGCs, and immature abDGCs in particular, in spatial cognition tasks that necessitate resolving memory interference ^8,16,28,39-42,71-74,19,32,35,75-77^. In sharp contrast, the role of abDGCs in social cognition is much less understood. One study showed that chemogenetic inhibition of mossy fiber terminals of a mixed population of immature abDGCs impaired remote memory retrieval underlying pup-mother recognition and emergent network properties^37^. Whether abDGCs contribute to other facets of social cognition such as encoding and consolidation of social recognition memory and resolution of social memory interference is not clear. Importantly, the neural circuit mechanisms by which abDGCs functionally modify network properties to support social recognition are not instantiated.

Circuit-based frameworks for conceptualizing the role of abDGCs in cognition must incorporate emerging insights from analyses of adult hippocampal neurogenesis in higher mammals. How neoteny of abDGCs affects cognition and underlying circuit mechanisms is poorly understood. Here, we engineered a genetic strategy to model the effective impact of neoteny of abDGCs on an inhibitory circuit mechanism that we identified as a substrate for social recognition^78,79^. We show that selective synchronous expansion of a single cohort of 4 weeks old immature, but not 8 weeks old mature abDGCs, increases functional recruitment of fast spiking parvalbumin expressing inhibitory interneurons (PV INs) in CA3/CA2, number of PV IN-CA2 and PV IN-CA3 synapses, mossy fiber-dependent recruitment of GABAergic inhibition of CA3 and CA2 and social memory consolidation in a pro-active memory interference task*. In vivo* local field potential recordings showed that the genetic expansion of a 4-week-old abDGC cohort enhances the baseline power, amplitude, and duration of sharp-wave ripples (SWRs) in CA1 and CA2, a neural substrate for memory consolidation. Additionally, it increased SWR rate sensitivity to social interaction. Chemogenetic manipulations targeting inhibitory neurons demonstrate that CA3/CA2 INs, including PV INs, are necessary for social memory consolidation after genetic expansion of a 4-week-old abDGC cohort and sufficient for this process in wild-type mice. Together, our findings suggest that the neoteny of abDGCs may be an evolutionary adaptation that enhances cognition by reshaping PV IN circuitry and modifying network properties essential for memory consolidation.

## Results

### Neotenic expansion of abDGC population decreases social memory interference

Neoteny of abDGCs maintains a small population of abDGCs in an immature state for a prolonged period of time. Therefore, we reasoned that the effective impact of neoteny of abDGCs on circuitry can be modeled in mice by integrating two variables, size of abDGC population and abDGC cell state. This can be done in two complementary ways. First, slow down the maturation of abDGCs in the adult DG (small number of abDGCs X extended window of immature cell state). Second, expand a single cohort of age-matched immature or mature abDGCs in adult DG (larger number of abDGCs X short window of immature cell state). Since the first approach is encumbered by risk of maladaptive integration of abDGCs, we devised a genetic strategy to selectively expand a single population of immature (4 weeks old) or mature (8 weeks old) abDGCs in adult DG of mice (**Fig.1a**). To this end, we generated adult Ascl1CreERT^2^: Bax ^f/f^ mice (iBax^Ascl1^) in which the pro-apoptotic gene *Bax* is conditionally recombined by tamoxifen administration in neural progenitors and activated neural stem cells of adult DG^80^. Inducible deletion of *Bax* promotes survival and synaptic integration of abDGCs^16,30^. This strategy contrasts with prior work using the Nestin CreERT^2^ line that targets *Bax* recombination in all neural stem cells and progenitors in adult DG thereby resulting in an expanded cohort of abDGCs of mixed ages ^30^. We bred iBax^Ascl1^mice with mice carrying the Cre sensitive reporter mouse line LSL-tdTomato (Ai14, B6;129S6-*Gt(ROSA)26Sor^tm14(CAG-tdTomato)Hze^*/J)^81^ to quantify the number of abDGCs following *Bax* recombination. At 4 weeks and 8 weeks post tamoxifen administration, iBax^Ascl1^:Ai14 mice exhibited a significant expansion of age-matched population of 4 weeks old or 8 weeks old abDGCs, respectively (**Fig. 1b**).

**Figure 1.**
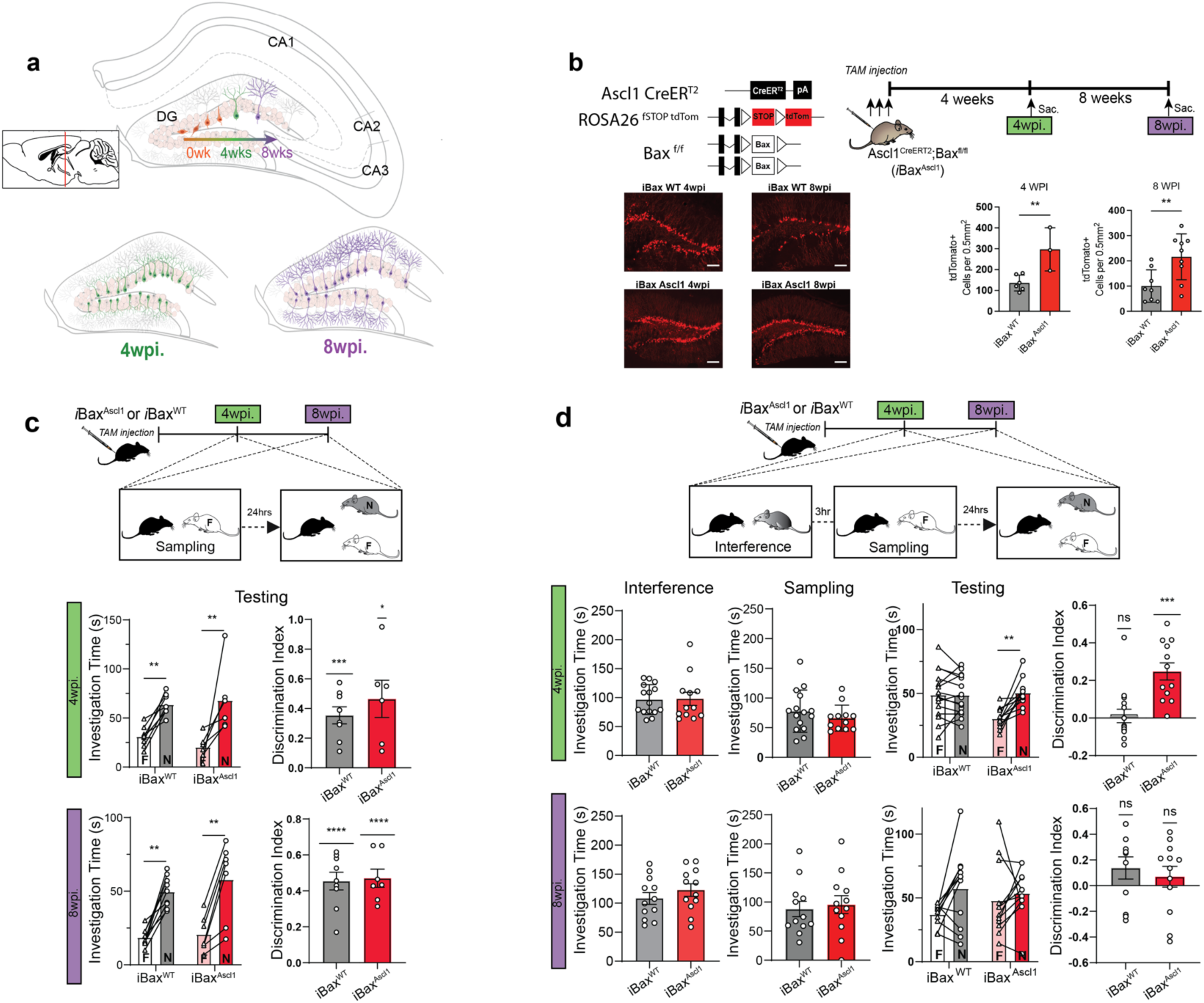
Genetic expansion of a 4 weeks old-abDGC population decreases social memory interference. **a,** Modelling effective impact of neoteny of abDGCs on hippocampal circuitry and function. Schematic illustrating how DG is modified by integration of a single expanded cohort of 4 weeks old or 8 weeks old abDGCs. **b,**Top. Schematic showing genetic strategy to selectively expand a single cohort of 4 weeks old- or 8 weeks-old abDGCs and use of Ai14 reporter to genetically label abDGCs. Bottom. Representative images of hippocampal sections obtained from 4 weeks group and 8 weeks group iBax^Ascl1 or WT^: Ai14 mice. Quantification of 4 weeks old or 8 weeks old tdTomato positive abDGCs reveals an approximate 2-fold increase in iBax^Ascl1^:Ai14 mice compared to iBax ^WT^: Ai14 mice. Scale bar = 100 µm **c,** 4 weeks group and 8 weeks group iBax^Ascl1^ mice (4 wpi.: n = 6; 8 wpi.: n = 7) exhibit comparable discrimination of novel vs. familiar social stimulus as iBax ^WT^ mice (4 wpi.: n = 8; 8 wpi.: n = 8) in SRM task. Two-way Repeated Measure ANOVA (genotype X Investigation time) with Šídák’s multiple comparisons test was used for investigation time comparisons. One-sample t-test were used for discrimination index comparisons between groups, \**p* < 0.05, \*\**p* < 0.005, \*\*\**p* < 0.001, \*\*\*\**p* < 0.0001. **d,** 4 weeks group but not 8 weeks group iBax^Ascl1^ mice exhibit discrimination of novel vs. familiar social stimulus in SRMi (SRM +pro-active interference) task. No difference in sampling in investigation time is observed between groups in interference and sampling phases. iBax^Ascl1^ mice (4 wpi.: n = 12; 8 wpi.: n = 12) iBax ^WT^ mice (4 wpi.: n = 15; 8 wpi.: n = 12) per group, Two-way Repeated Measure ANOVA (genotype X Investigation time) with Šídák’s multiple comparisons test was used for investigation time comparisons. One-sample t-test were used for discrimination index comparisons between groups, \*\**p* < 0.005, \*\*\**p* < 0.001.

Next, we tested whether neotenic expansion (mice with expanded cohort of 4 weeks old abDGCs) enhances social recognition memory (SRM). In the SRM task, a subject mouse is challenged to distinguish between a familiar social stimulus encountered 24 hours previously and a novel social stimulus. Like their wild-type littermates (*Bax^f/f^*or iBax^WT^ mice), both 4 week- and 8 week-groups of iBax^Ascl1^ mice significantly discriminated between the familiar and novel mouse (**Fig. 1c**). SRM is highly sensitive to proactive interference, whereby recently acquired, overlapping social memories affect social memory encoding and consolidation^82^. We modified the SRM task so that the subject mouse encounters a social stimulus three hours prior to encoding the familiar social stimulus as shown in **Fig. 1d**. In this SRM interference or SRMi task, the subject mouse must resolve pro-active interference to encode the social stimulus in sampling phase and consolidate the familiar social stimulus in order to successfully discriminate between the familiar and novel mouse. In the SRMi task, only the 4 week group of iBax^Ascl1^ mice but not the 8 week group of iBax^Ascl1^ mice or iBax^WT^ mice successfully discriminated between the familiar and novel mouse (**Fig. 1d**). All groups of mice exhibited equivalent levels of investigation of interference stimulus and familiar stimulus during sampling phase (**Fig. 1d**). Together, these observations suggest that neotenic expansion of abDGC population enhances social recognition memory consolidation in a behavioral interference task.

Neurogenesis-dependent forgetting has been proposed as a mechanism to reduce proactive memory interference^44^. To determine whether forgetting of the proactive stimulus is a putative mechanism by which memory interference is decreased in 4 week group iBax^Ascl1^ mice, we tested iBax^Ascl1^ mice in the contextual fear conditioning paradigm. This task is ideal for assessing how increasing neurogenesis following learning affects memory strength, an indicator of forgetting^83^. Expansion of a 4 week immature abDGC population following contextual fear learning did not affect the strength of fear memory tested four weeks later, consistent with prior results^16^ arguing against a role for neurogenesis-mediated forgetting in iBax^Ascl1^ mice (**Extended Data Fig. 1a, b**).

### Neotenic expansion of abDGC population increases PV IN synapses, basal inhibitory synaptic transmission and feed-forward inhibition in DG-CA3/CA2

Previously, we showed that PV INs in response to increased mossy fiber excitatory drive increased their perisomatic inhibitory synapses onto CA3 and CA2 pyramidal neurons ^78,79^, a hub for social memory^62,84-90^. Additionally, we implicated PV IN mediated feed-forward inhibition in DG-CA3 and DG-CA2 as a neural circuit mechanism for social recognition ^79,78^. To understand how expansion of immature and mature abDGCs affects PV IN-mediated FFI in DG-CA3 and DG-CA2, we first quantified PV IN perisomatic contacts onto CA3/CA2 subregions 4- and 8-weeks post tamoxifen induction (**Fig.2a**). At the 4-week but not 8-week timepoint, iBax^Ascl1^ mice exhibited significantly higher number of PV IN synapses (PV Puncta) in CA3 and CA2 than iBax^WT^ mice (**Fig.2b,c**). Reflecting these anatomical changes in PV IN-CA3/CA2 pyramidal neuron connectivity, *ex vivo* whole-cell recordings of CA3/CA2 pyramidal neurons revealed an increase in the frequency of miniature inhibitory postsynaptic current (mIPSC) 4 weeks post induction (**Fig. 2d,e,f**). There were no significant differences in the amplitude between groups at the 4-week timepoint and no differences in these measures at 8 weeks post induction (**Fig. 2e, f**). Recordings from PV INs along the stratum lucidum revealed an increase in the frequency of miniature excitatory postsynaptic current (mEPSC) in iBax^Ascl1^ mice 4 weeks post induction (**Fig. 2g**). Additionally, there were no differences in mEPSC recordings from CA3/CA2 pyramidal neurons between groups at either 4- or 8-weeks post induction (**Extended Data Fig. 2a, b**). These data suggest that expanding the immature, but not mature, abDGC population increases excitatory transmission onto PV INs, PV IN synapses in CA3/CA2 and basal inhibitory synaptic transmission onto CA3/CA2 pyramidal neurons.

**Figure 2.**
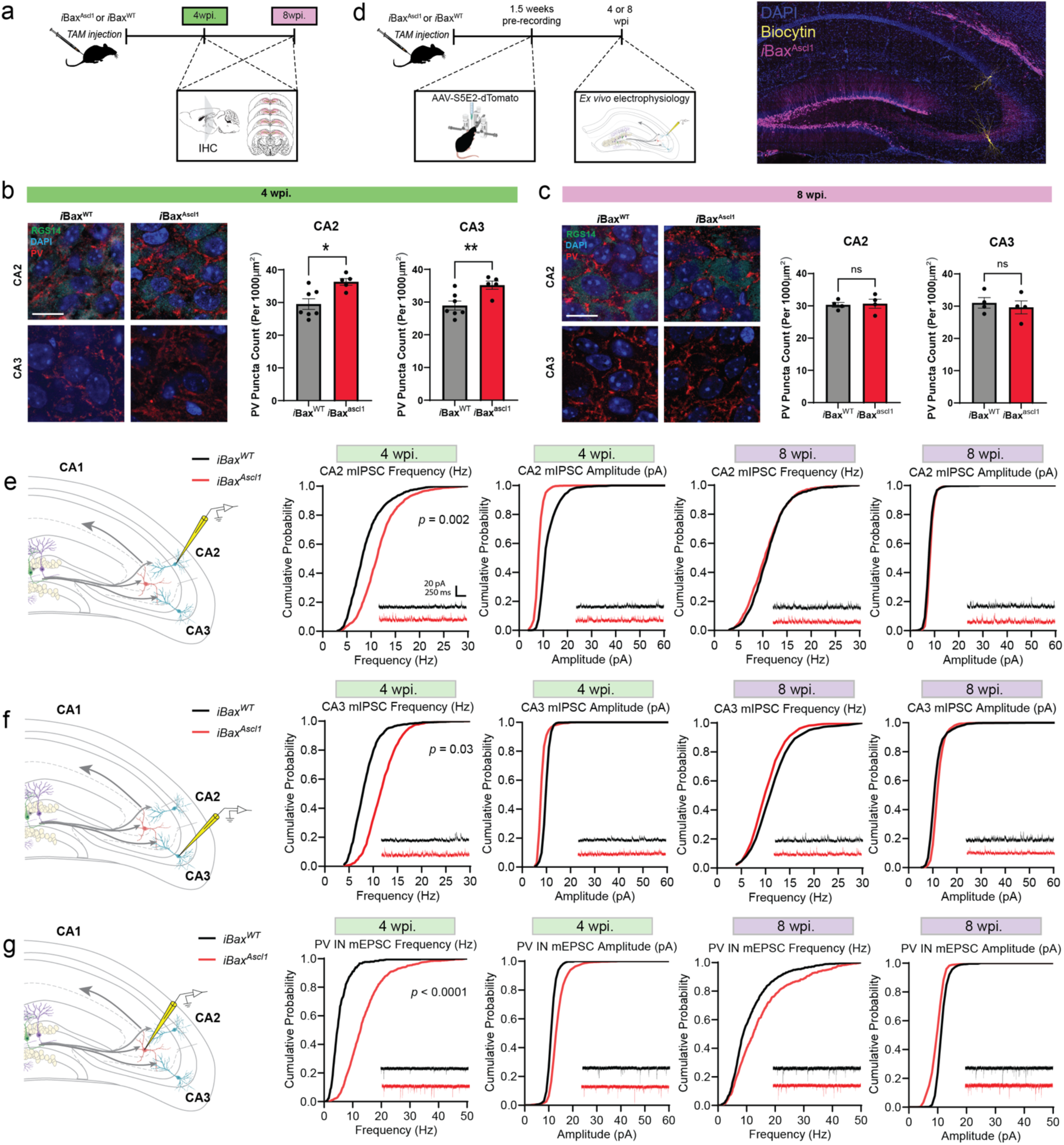
Expansion of single cohort of 4 weeks-, but not 8 weeks-old abDGCs, increases basal inhibitory synaptic transmission in CA3 and CA2. **a,** Experimental design. PV immunohistochemistry (IHC) was performed 4- and 8-weeks post TAM. **b,** Representative images and quantification of PV^+^ puncta density in CA2/CA3 stratum pyramidale 4 weeks post TAM injection. RGS14^+^ labeling defines CA2 subfield. Two-tailed unpaired *t* test was used between groups, n = 5-7 mice per group, \**p* < 0.05, Mean ± SD, scale bar = 15 µm. **c,** Representative images and quantification of PV^+^ puncta density in CA2/CA3 stratum pyramidale 8 weeks post TAM injection. RGS14^+^ labeling defines CA2 subfield. Two-tailed unpaired *t* test was used between groups, n = 4 mice per group. Mean ± SD, scale bar = 15 µm. **d,** Left. Schematic depicting AAV-S5E2-dTom injected into dorsal CA3/CA2 1.5 weeks prior to ex vivo recordings from 4- or 8-week post TAM injection. Right. Representative image of dorsal hippocampal section with genetic expansion of abDGC population. Biocytin filled neurons were patched in CA3 and CA2. **e,** Schematic depicting whole-cell patch-clamp recording of miniature inhibitory postsynaptic current (mIPSC) from CA2 pyramidal neurons (PN) (left). Cumulative probability plots of mIPSC frequency and amplitude from PNs in 4- and 8-week post TAM injection. Kolmogorov-Smirnov test was used between groups, n = 11-14 cells, 2-4 cells per mouse, 4 mice per group. **f,** Schematic depicting whole-cell patch-clamp recording of mIPSC from CA3 PNs (left). Cumulative probability plots of mIPSC frequency and amplitude from PNs in 4- and 8-week post TAM injection. Kolmogorov-Smirnov test was used between groups, n = 9-14 cells, 2-4 cells per mouse, 3-4 mice per group. **g,** Schematic depicting whole-cell patch-clamp recording of miniature excitatory postsynaptic current (mEPSC) from stratum lucidum parvalbumin^+^ interneurons (PV IN) (left). Cumulative probability plots of mEPSC frequency and amplitude from PV INs in 4- and 8-week post TAM injection. Kolmogorov-Smirnov test was used between groups, n = 15-18 cells, 4-7 cells per mouse, 3-4 mice per group.

We next asked how immature or mature abDGC expansion affects mossy fiber recruitment of feed-forward inhibition of CA3 and CA2. We expressed Channelrhodopsin (ChR2) 1.5 weeks before whole-cell recordings from pyramidal neurons in 4 and 8 weeks adult iBax^Ascl1^ mice and iBax^WT^ littermates (**Fig. 3a,b**). Analysis of optically evoked, mossy-fiber driven excitation upon CA2 pyramidal neurons revealed that in 4 weeks-but not 8 weeks-iBax^Ascl1^ mice there was an increase in inhibition in proportion to excitation (**Fig. 3c,d**). There was a significantly higher IPSC paired pulse ratio in iBax^Ascl1^ mice compared to controls at 4 weeks post induction which was due to both IPSC responses having a high IPSC amplitude (**Fig. 3d, Extended Data Fig. 3a**). We note that iBax^Ascl1^ mice had a reduced paired pulse ratio of the IPSC from CA2 pyramidal neurons at the 8-week timepoint (**Extended Data Fig. 3a**). This indicates a higher vesicle release probability of inhibition, however the IPSC amplitude was not statistically different than controls (**Fig. 3d**). We observed similar trends in CA3 pyramidal neurons, however there were no statistically significant differences between groups across these measures (**Fig. 3e, Extended Data Fig. 3b**). Together, these findings demonstrate that neotenic expansion of an abDGC population increases mossy fiber driven feed-forward inhibition of CA2. Since mossy fibers of abDGCs in 4 weeks-iBax^Ascl1^ mice represent a small fraction of mossy fibers expressing ChR2 in these experiments, our data convey how immature abDGCs disproportionately recruit feed-forward inhibition of CA2.

**Figure 3.**
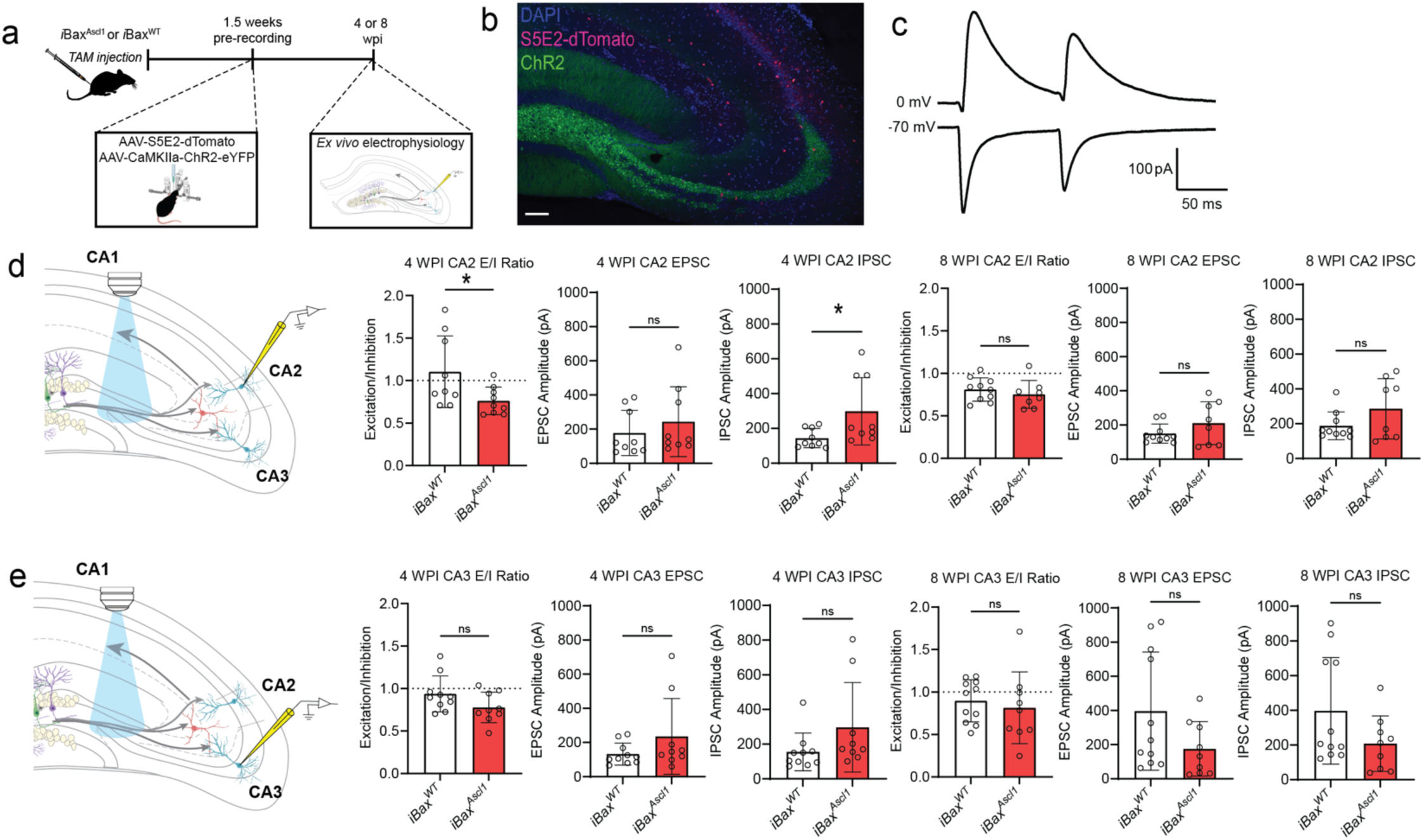
Expansion of single cohort of 4 weeks-, but not 8 weeks-old abDGCs, increases feed-forward inhibition of CA2. **a,** Schematic depicting AAV-S5E2-dTomato injected into dorsal CA3/CA2 and rAAV5-CaMKIIα-ChR2-eYFP injected into dorsal dentate gyrus (DG) 1.5 weeks prior to ex vivo recordings from 4- or 8-week post TAM injection. **b,** Representative image of dorsal hippocampal section with genetic expansion of abDGC expressing ChR2-eYFP and PV targeted S5E2-dTomato. Scale bar = 100 µm. **c,** Representative *ex vivo* traces depicting EPSC and IPSC responses to optically evoked (473 nm) paired pulse stimulation. **d,** Schematic depicting whole-cell patch-clamp recording of optically evoked EPSC and IPSC from CA2 PNs (left). Bar graphs depict excitation to inhibition (E/I) ratio and the amplitude of the first EPSC and IPSC response to paired pulse optical stimulation. Two-tailed unpaired *t* test was used between groups, n = 9-10 cells, 2-4 cells per mouse, 3-4 mice per group, \**p* < 0.05, Mean ± SD. **e,** Schematic depicting whole-cell patch-clamp recording of optically evoked EPSC and IPSC from CA3 PNs (left). Bar graphs depict excitation to inhibition (E/I) ratio and the amplitude of the first EPSC and IPSC response to paired pulse optical stimulation. Two-tailed unpaired *t* test was used between groups, n = 9-11 cells, 2-4 cells per mouse, 3-4 mice per group, Mean ± SD.

### Chemogenetic activation of CA3/CA2 PV INs is sufficient to decrease social memory interference

We next asked whether chemogenetic activation of PV INs is sufficient to decrease social memory interference. We expressed the chemogenetic activator hM3D(Gq)-DREADD-dTom ^91^ or dTom in CA3/CA2 PV INs of adult iBax^WT^ mice using PV IN-enhancer driven viral vectors ^78,79,92^ and administered Clozapine-N-oxide (CNO) one hour prior to sampling phase (**Fig. 4a, b**). As reported earlier in **Fig. 1d**, iBax^WT^ mice failed to discriminate between the familiar and novel social stimulus (**Fig. 4c)**. In contrast, chemogenetic activation of CA3/CA2 PV INs in iBax^WT^ mice resulted in significant discrimination between the familiar and novel social stimulus (**Fig. 4c)**. Both groups of mice exhibited comparable levels of investigation of social stimulus in interference and sampling phases suggesting that PV IN activation enhanced memory consolidation. Given the half-life of CNO^91^, the reduction in memory interference is likely to reflect enhanced encoding and consolidation of the familiar stimulus in the SRMi task.

**Figure 4.**
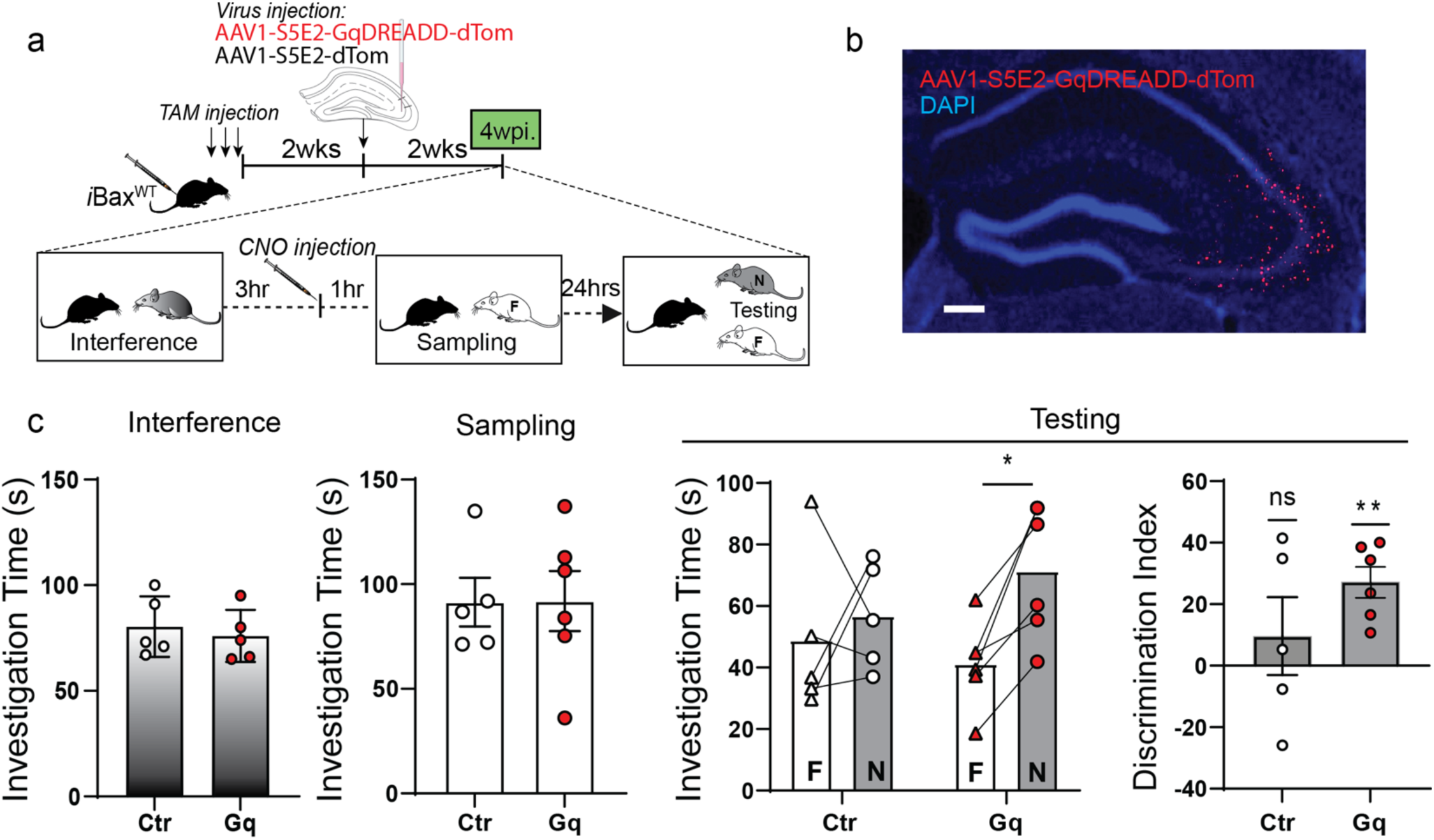
Chemogenetic activation of CA3/CA2 PV INs is sufficient to decrease social memory interference. **a,** Experimental design: iBax^WT^ mice were tested in SRMi task two weeks after PV targeted viral injection of AAV-S5E2-hM3D(Gq)-DREADD-dTom in CA2/3. Wild-type mice received TAM injections to maintain consistency of protocol and TAM exposure across experiments. **b,** Representative image of dorsal hippocampus section showing viral expression of S5E2-hM3D(Gq)-DREADD-dTom in CA2/3. Scale bar = 15 µm. **c,** Chemogenetic activation of CA2/CA3 PV INs before sampling significantly improved discrimination of social stimuli during testing in SRMi task (*p<0.05). No difference in social stimulus investigation during interference or sampling sessions was observed. Two-way Repeated Measure ANOVA (genotype X Investigation time) with Šídák’s multiple comparisons test was used for investigation time comparisons. One-sample t-test were used for discrimination index comparisons (control virus: n = 5, Gq-DREADD n = 6), \**p* < 0.05, \*\**p* < 0.005.

### Chemogenetic inhibition of CA3/CA2 INs abrogates the effect of neotenic expansion of abDGC population on social memory interference

We next asked if intact inhibition of CA3/CA2 in 4 weeks- iBax^Ascl1^ mice is necessary for social memory consolidation in SRMi task. We expressed the hM4Di(Gi)-DREADD-dTom or dTom in CA3/CA2 INs of adult 4 weeks- iBax^Ascl1^ mice using a Dlx 1/2 enhancer driven viral vector ^93^. CA3/CA2 INs were inactivated by administration of systemic CNO 1 hour prior to sampling phase (**Fig. 5a)**. Unlike the control group of 4 weeks- iBax^Ascl1^ mice, 4 weeks- iBax^Ascl1^ mice in which CA3/CA2 INs were silenced failed to discriminate between the familiar and novel social stimulus (**Fig. 5b)**. Chemogenetic silencing of CA3/CA2 INs in 4 weeks- iBax^Ascl1^ mice 1 hour prior to testing impaired discrimination between the familiar and novel social stimulus suggesting a role for these INs in memory retrieval (**Fig. 5c,d)**. Together, these results suggest that CA3/CA2 INs are necessary for mediating the resolving effects of neotenic expansion of an abDGC population on social memory interference.

**Figure 5.**
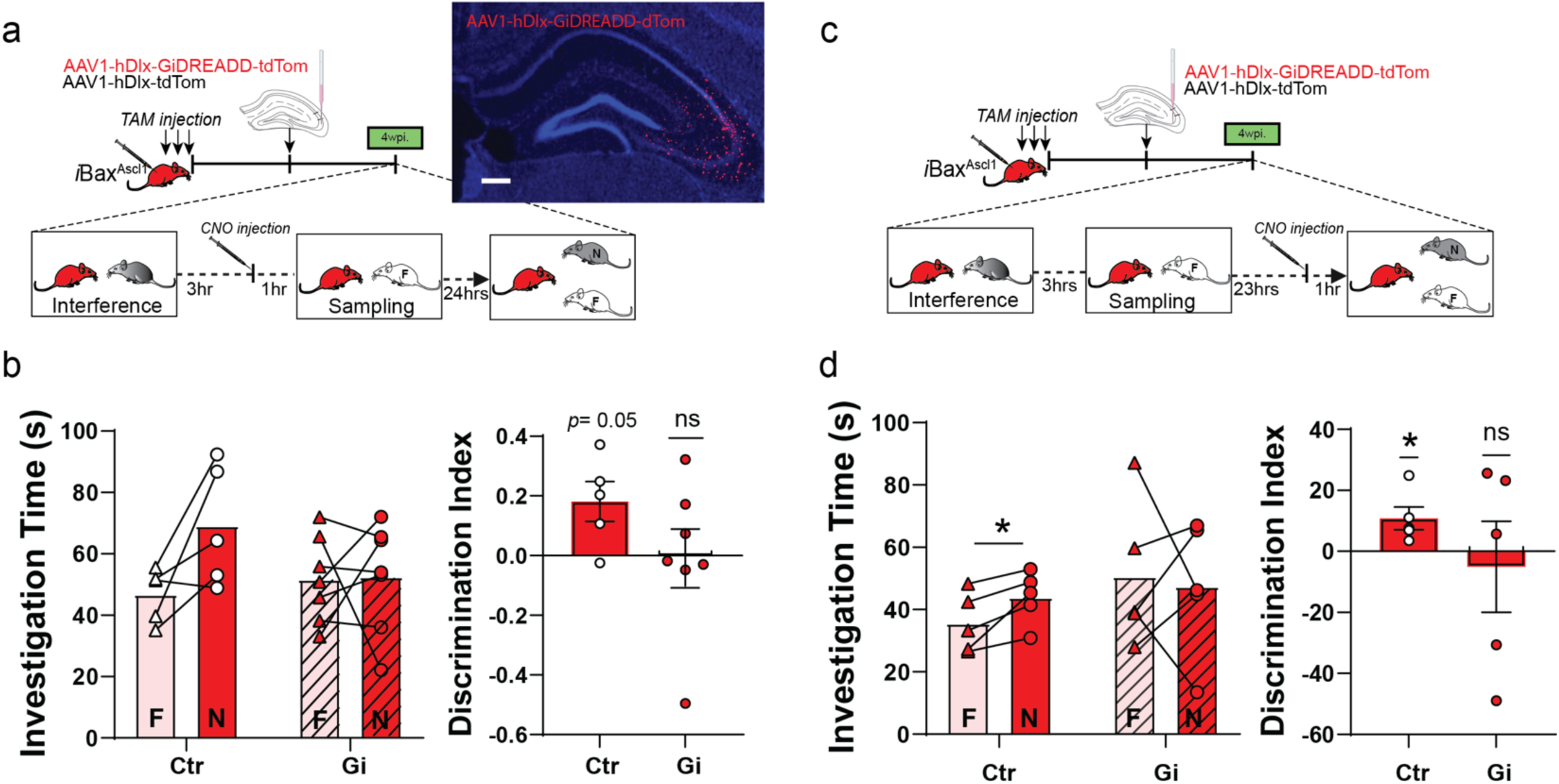
Chemogenetic inhibition of CA3/CA2 INs abolishes the effect of neotenic expansion of abDGC population on social memory interference. **a,** Left. Experimental design: iBax^Ascl1^ mice were tested in SRMi task 4 weeks post TAM and two weeks after viral injection of AAV-Dlx1/2-hM4Di(Gi)-DREADD-dTom or control AAV-Dlx1/2-dTom in CA2/CA3. CNO was administered 1 hour prior to sampling phase. Right. Representative image of dorsal hippocampus section showing viral expression of hM4Di(Gi)-DREADD-dTom in CA2/CA3. Scale bar = 15 µm. **b,** Chemogenetic inactivation of CA2/CA3 INs prior to sampling phase impaired discrimination between novel and familiar mice. The control group (iBax^Ascl1^ mice with control AAV-Dlx1/2-dTom virus) successfully discriminated between social stimuli. Two-way Repeated Measure ANOVA (genotype X Investigation time) and paired t-test were used for investigation time comparisons. One-sample t-test were used for discrimination index comparisons between groups (control virus: n = 5, Gi-DREADD n = 7), \**p* < 0.05. **c,** Experimental design: iBax^Ascl1^ mice were tested in SRMi task 4 weeks post TAM and two weeks after viral injection of AAV-Dlx1/2-hM4Di(Gi)-DREADD-dTom or control AAV-Dlx1/2-dTom in CA2/CA3. CNO was administered 1 hour prior to testing phase. **d,** Chemogenetic inactivation of CA2/CA3 INs prior to sampling phase impaired discrimination between novel and familiar mice. The control group (iBax^Ascl1^ mice with control AAV-Dlx1/2-dTom virus) successfully discriminated between social stimuli. Two-way Repeated Measure ANOVA (genotype X Investigation time) and paired t-test were used for investigation time comparisons. One-sample t-test were used for discrimination index comparisons (control virus: n = 5, Gi-DREADD n = 5), \**p* < 0.05.

### Neotenic expansion of abDGC population increases baseline SWR properties and enhances rate change sensitivity to social interaction in CA1 and CA2

PV IN mediated perisomatic inhibition of principal cells is necessary for synchronizing principal cell activity to generate SWRs ^94-97^, a neural substrate for memory consolidation^87,98-102^. Optogenetically extending ripple length was shown to result in enhanced spatial memory^101^. In prior work, we found that selectively increasing feed-forward inhibition in DG-CA3/CA2 results in reconfiguration of network properties such as SWR-spindle coupling to support memory consolidation^103^. Since genetic expansion of immature abDGCs results in increased recruitment of PV INs and feed-forward inhibition of CA2 and CA3, we asked how SWR properties change in CA2 and CA1 of 4 weeks- *i*Bax^Ascl1^ mice. Multi-channel electrodes targeting CA1, CA2, and the DG were chronically implanted in iBax^Ascl1^ and iBax^WT^ mice. After 1 week of full recovery from the implant surgeries, baseline local field potentials (LFP) were recorded during 2 hour sleep sessions in home cages (0-week timepoint). Following baseline measurements, LFP recordings were performed during 2 hour sleep sessions at 4 and 8 weeks post tamoxifen injections within the same mice (**Fig. 6a-d**). In contrast to iBax^WT^ mice, 4 weeks- iBax^Ascl1^ mice exhibited a significant increase in baseline CA1 ripple power, amplitude, duration and rate (0 vs. 4wpi) (**Fig. 6e, f**). In CA2, 4 weeks- iBax^Ascl1^ mice exhibited a significant increase in baseline ripple power, amplitude and duration (0 vs. 4wpi). No further changes were observed at the 8 week timepoint (**Fig. 6e, g**). The persistent change in some ripple properties could be due to re-exposure to social stimulus at 4 week timepoint. We also tested whether social experience during SRMi task changes CA1/CA2 ripple properties by measuring sleep LFPs between each of the sessions (interference, sampling and testing) at 4 weeks post tamoxifen induction. Consistent with prior work^87^, we observed significant changes in CA1 and CA2 ripple rates in iBax^Ascl1^ mice following social stimulus investigation in both the sampling and interference phases of the SRMi task **(Fig. 7 a-c).** Notably, the iBax^Ascl1^ group showed a significant increase in the CA1 ripple rate after two consecutive social interactions spaced three hours apart whereas iBax^WT^ mice exhibited an increase in the CA1 ripple rate only after the very first social interaction **(Fig. 7 b,c)**. Additionally, iBax^Ascl1^ group showed significantly enhanced ripple rate after the first social interaction (**Fig. 7c**). There were no significant changes in ripple properties (power and amplitude) except that iBax^WT^ mice showed a significant increase in CA1 ripple duration after the first exposure to the first animal **(Extended Data Fig.4)**. The relatively small increase in CA2 ripple rate in iBax^WT^ after social interaction may reflect the loss of social stimulus-associated novelty because our mice were group housed to prevent detrimental effects of social isolation on adult hippocampal neurogenesis^104^. However, the iBax^Ascl1^ mice still exhibited heightened sensitivity of CA1/2 ripple rate to social interaction. Together, these data suggest that neotenic expansion of the abDGC population modifies CA2 and CA1 SWR properties into a configuration (increased power, duration, and rate) that is permissive for social memory encoding and consolidation^87,97,101^.

**Figure 6.**
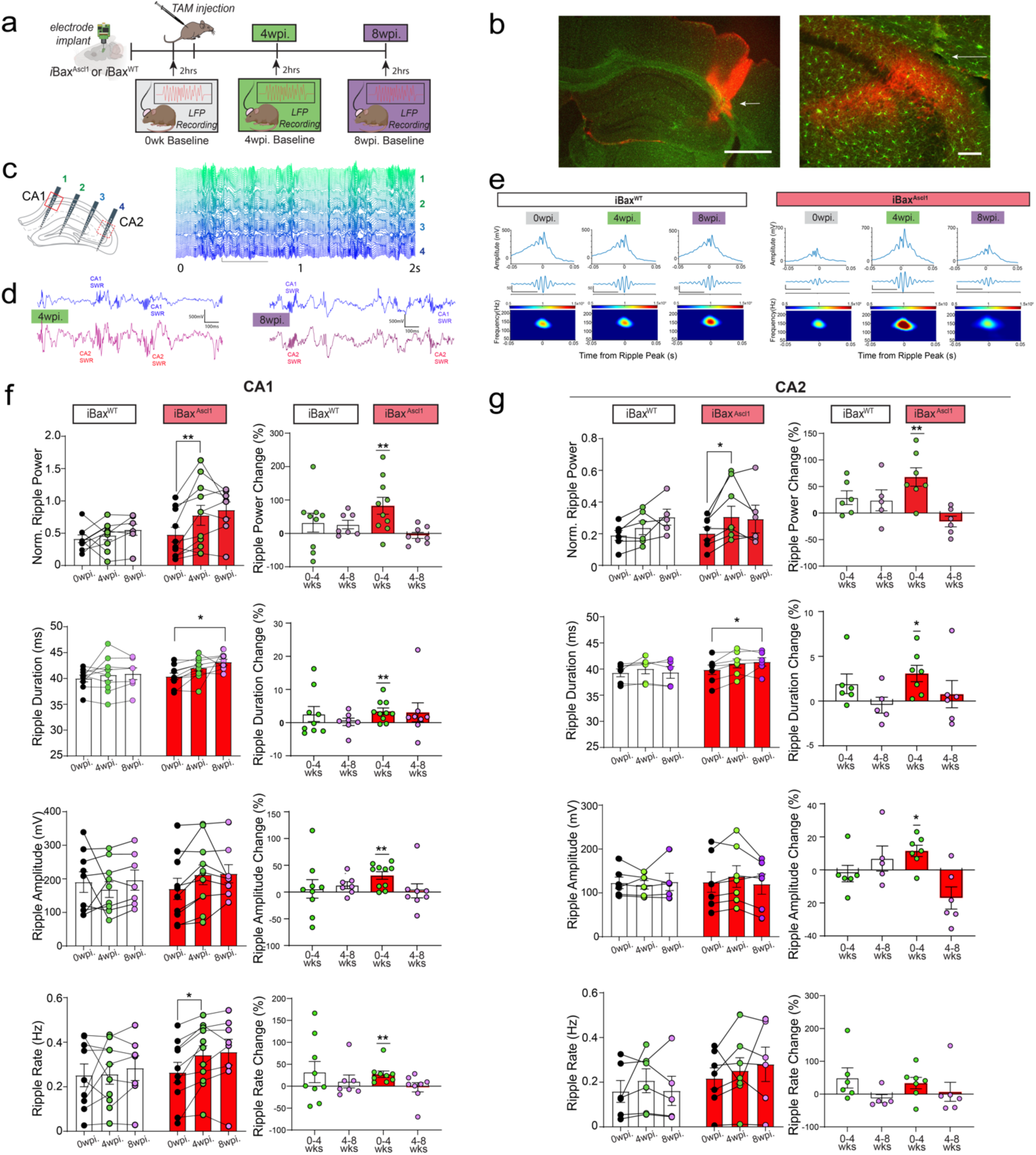
Neotenic expansion of abDGC population increases power, amplitude and duration of SWRs in CA1 and CA2. **a,** Experimental schedules: Electrodes were chronically implanted in iBax^WT^ and iBax^Ascl1^ mice. Baseline local field potential (LFP) signals were measured for 2 hours during sleep after recovery. At 4- and 8-weeks following TAM injection, LFP signals were recorded during 2 hour sleep sessions. **b,** Left. A histology figure indicating electrode locations with Dil dye (red) (the location of the shank targeting CA2 is marked with a white arrow). Scale bar = 1mm. Right. RGS14 (red) immunohistochemistry image confirmed CA2 electrode locations (white arrow). Scale bar = 15 µm. **c,** Left. Subset of mice were implanted with 64 channel multi-shank probes that span all the layers of hippocampus including CA1 and CA2 to measure CA1 and CA2 ripples simultaneously. Right. Two seconds of 64 channel example traces of hippocampus LFP during NREM sleep. Scale bar = 50 µV/400ms. **d,** Example traces of simultaneous CA1 and CA2 ripple recordings in the same mouse shows stable ripple signaling in CA1 and CA2 at 4- and 8-weeks following TAM injection. Scale bar = 500 µV/100ms. **e**, Left. Averaged ripple trace from a single iBax^WT^ mouse at 0, 4, and 8 weeks post-TAM showed stable recordings (left). In contrast, the averaged ripple trace from an iBax^Ascl1^ mouse showed increased ripple signals at 4 weeks after TAM injection that decreased back to a level similar to the 0-week baseline (right). Scale bar = 50 µV/400ms **f**, Left column: CA1 ripple analysis showed significant increase in ripple power and rate from the 0-week to the 4-week time point in the iBax^Ascl1^ group. Normalized ripple power was defined as the ratio of ripple power to delta power during all NREM epochs. Right column: Percentage changes showed a significant increase in power, duration, amplitude, and rate. iBax^WT^ mice did not show any significant changes over time. 10-11 mice per group. Two-way Repeated Measure ANOVA (genotype X wpi.) with Šídák’s multiple comparisons test was used for post hoc group comparisons. One-sample t-test were used for percentage change comparisons comparisons for 0- and 4-week post injections or 4- and 8 week post injections, \**p* < 0.05, \*\**p* < 0.005. **g**, Left column: CA2 ripple analysis showed significant increase in ripple power and duration from the 0-week to the 4-week time point in the iBax^Ascl1^ group. Right column: Percentage changes also showed significant increases in ripple power, amplitude and duration from the 0-week to the 4-week time point in the iBax^Ascl1^ group. iBax^WT^ mice did not show any significant changes over time. 5-7 mice per group. Two-way Repeated Measure ANOVA (genotype X wpi.) with Šídák’s multiple comparisons test was used for post hoc group comparisons. One-sample t-test were used for percentage change comparisons for 0- and 4-week post injections or 4- and 8 week post injections, \**p* < 0.05, \*\**p* < 0.005.

**Figure 7.**
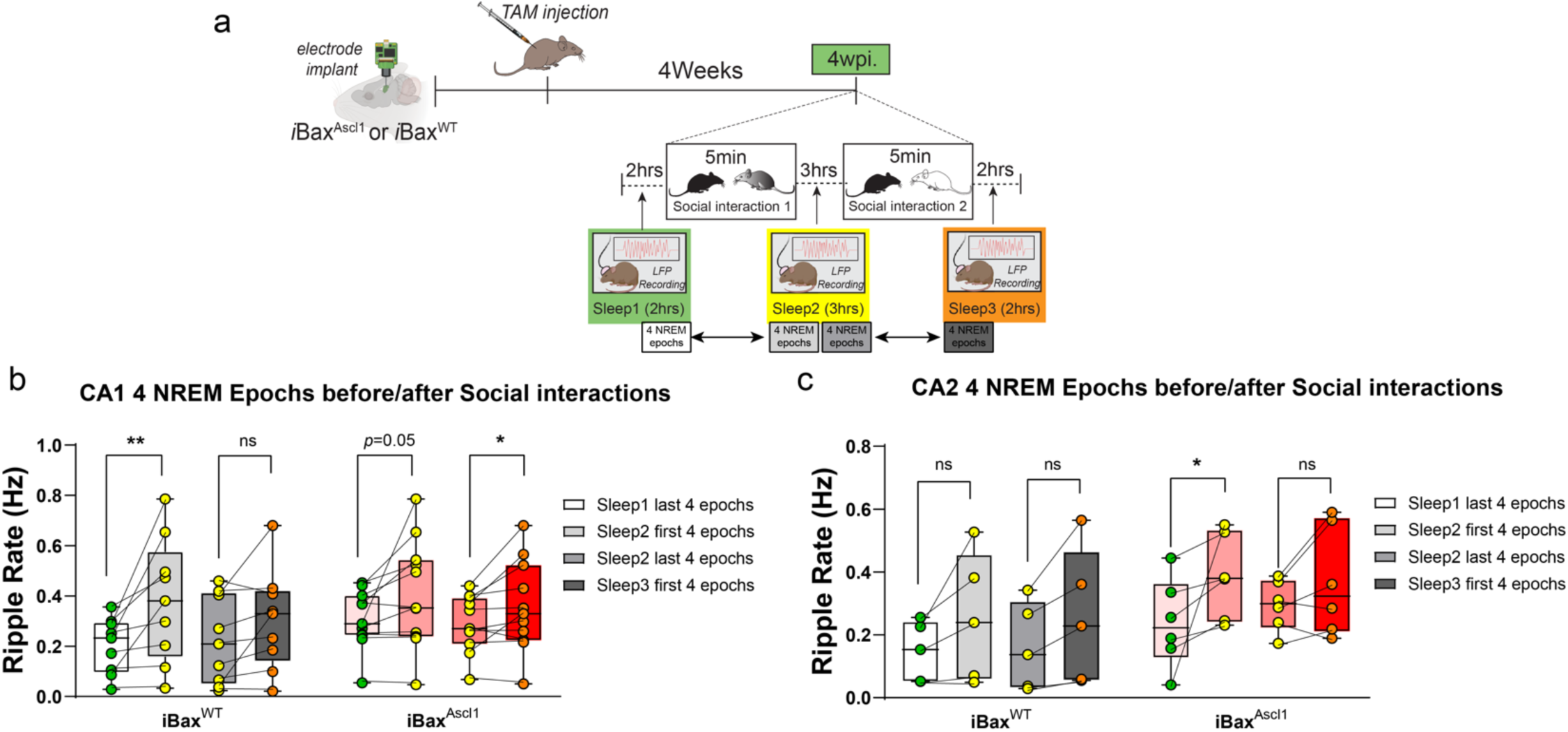
Social experience significantly increased ripple rate. **a,** Experimental schedules: Electrodes were chronically implanted in iBax^WT^ and iBax^Ascl1^ mice. After recovery, TAM was injected in both iBax^WT^ and iBax^Ascl1^ groups. 4 weeks after TAM injection baseline local field potential (LFP) signals were measured for 2 hours during sleep (Sleep 1). After 5 minutes of the first social interaction, all mice underwent 3 hours of home cage sleep sessions (Sleep 2). This was followed by another 5-minute social interaction with a novel mouse and a subsequent 2-hour sleep session (Sleep 3). **b,** After the first 5-minute exposure to the novel mice, both the iBax^WT^ and iBax^Ascl1^ groups showed a significant increase of CA1 ripple rate. However, only the iBax^Ascl1^ group showed a significant ripple rate increase after the second exposure to the novel juvenile mouse (iBax^WT^ : n= 9, iBax^Ascl1^ : n = 11), Paired t-test was used for before-after social exposure ripple rate changes. \**p* < 0.05. **c,** In the iBax^Ascl1^group, the CA2 ripple rate was significantly increased after the first social interaction (\**p* < 0.05). The second exposure to the novel mouse did not increase the CA2 ripple rate in either group(iBax^WT^ : n= 5, iBax^Ascl1^ : n = 6) Wilcoxon matched-pairs signed rank test was used for before-after social exposure ripple rate changes.

## Discussion

A significant body of work supports a role for abDGCs in decreasing spatial memory interference ^8,16,19,28,32,35,39-42,71-77^. In contrast, much less is known about how abDGCs contribute to social cognition. Evidence from analyses of adult hippocampal neurogenesis in non-human primates^48,49^ and to a lesser extent in humans has suggested that abDGCs exhibit neoteny^50-55^. Neoteny of abDGCs is hypothesized to offset decline in adult hippocampal neurogenesis by maintaining a population of abDGCs in an immature state when abDGCs exhibit unique physiological properties important for hippocampal-dependent memory operations^8^. However, empirical evidence is lacking. In this study we sought to address these two gaps in our understanding of how abDGCs contribute to cognition. To begin to test this thesis, we modelled neoteny of abDGCs by engineering mice in which we expanded a single population of 4 weeks old immature abDGCs or 8 weeks old mature abDGCs in the adult DG. Using these mice, we investigated whether neotenic expansion of abDGC population (4 weeks old abDGCs) improves social cognition. We found that expansion of a population of immature, but not mature abDGCs, enhanced consolidation of social recognition memory. This advantage conferred by neotenic expansion of abDGCs was manifest only when mice had to resolve pro-active social interference consistent with the role of the DG in decreasing memory interference^57-70^.

What are the neural circuit mechanisms by which neotenic expansion of abDGCs enhances social recognition? We found that modifying the DG with a single expanded cohort of 4 weeks-old abDGCs, but not 8 weeks-old abDGCs, resulted in increased excitatory drive onto PV INs, increased PV IN synapses in CA3/CA2 and feed-forward inhibition in DG-CA3/CA2. One mechanism by which a small number of immature abDGCs in iBax^Ascl1^ mice make a disproportionate contribution to mossy fiber-evoked inhibition of principal cells is through mossy fiber terminal-filopodial synapses onto PV INs ^105-108^, a learning-^107,109^ and social experience-sensitive synaptic substrate of feed-forward inhibition in DG-CA3/CA2^79,109^. Immature abDGCs exhibit higher mossy fiber terminal-filopodia than mature abDGCs ^20^(Guo and Sahay, unpublished observations). Furthermore, increased mossy fiber drive onto PV INs triggers a cell intrinsic plasticity program to increase and reorganize PV IN mediated perisomatic inhibition of downstream CA3/CA2 principal cells^78,79^. The combination of increased excitatory drive onto PV INs and resultant recruitment of PV IN-mediated perisomatic inhibition amplifies the contribution of a small number of immature abDGCs to modulation of hippocampal network activity^8,25,26,29,110^ (**Fig. 8**). Since feed-forward inhibition dictates spiking fidelity of principal cells and expands dynamic range of principal cell activity^111,112^, such a circuit mechanism may enable immature abDGCs to impose a sparse coding regimen, increase CA3/CA2 population dimensionality and consequently, enhance discrimination^35^. That chemogenetic activation of PV INs in wild-type mice was sufficient to enhance social memory consolidation in the SRMi task further implicates PV INs as critical arbiters of immature abDGC-dependent regulation of CA3/CA2 principal cell activity. Taken together, our findings define a neural circuit mechanism by which immature abDGCs modulate CA3/CA2 activity to support hippocampal dependent memory processing^8,25,26,29,110^.

**Figure 8.**
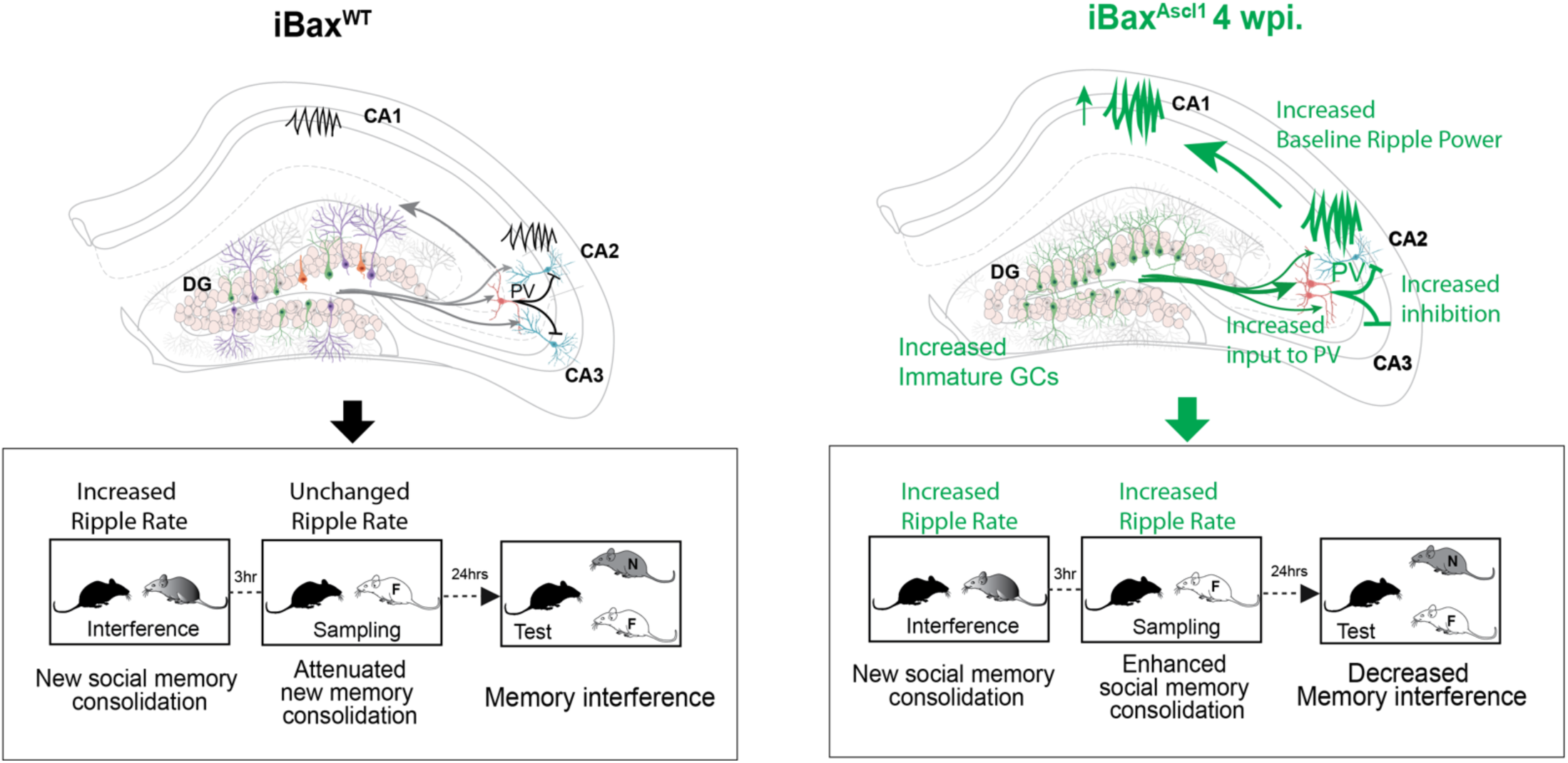
Neotenic expansion of an abDGC cohort decreases social memory interference by enhancing feed-forward inhibition and modifying CA1 and CA2 SWR properties. Genetic expansion of a single cohort of 4 weeks old abDGCs (iBax^Ascl1^ 4wpi mice, right) improved ability to recognize a novel animal during the test session of SRMi test compared to wild-type littermates (left). Genetic expansion of a single cohort of 4 weeks old abDGCs resulted in stronger PV IN-mediated feed-forward inhibition of CA2 and CA3 pyramidal neurons and increased CA1/CA2 ripple power and duration to decrease pro-active social memory interference and enhance social memory consolidation.

In contrast to prior reports ^44,113^, genetic expansion of age-matched abDGCs (this study) or genetic enhancement of neurogenesis ^16^ following learning does not promote forgetting. One way to resolve this discrepancy is to ascertain the extent to which the neural circuit mechanisms engaged by the genetic strategy used here and previously ^16,25,30^ are also engaged by genetic manipulations shown to promote forgetting.

Chemogenetic inhibition of PV INs in iBax^Ascl1^ mice in SRMi suggests a role for immature abDGC recruitment of PV INs in social memory consolidation. At a network level, PV INs are thought to be crucial for generation of SWRs through synchronous activity of principal cells^94-97^, and prolongation of SWR duration was shown to causally enhance spatial memory^101^. SWRs are thought to result from reciprocal interactions between PV INs, other INs and principal cells although the precise mechanisms are not resolved. Clues to potential mechanisms in CA3/CA2 come from several recent studies. Increasing feed-forward inhibition in DG-CA3/CA2 enhanced SWR-spindle coupling^103^. *In vivo* functional imaging of molecularly defined INs in CA3/CA2 identified unique activity profiles and suggested distinct contributions of PV INs (PV basket cells and axo-axonic cells) and Cholecystokinin interneurons (CCK INs) to SWR properties during learning^97^. Chemogenetic inhibition of mossy fiber terminals of immature abDGCs during social interaction impairs SWRs in awake mice ^37^. Our findings identify abDGC-recruitment of PV INs and feed-forward inhibition in DG-CA3/CA2 as a circuit mechanism by which immature abDGCs modify SWR properties in CA2 and CA1 during sleep to enhance social memory consolidation (**Fig. 8**). In this context, how abDGC recruitment of feed-forward inhibition affects burst firing of athorny CA3 pyramidal neurons, a trigger for SWRs, is not clear since these CA3 neurons do not receive mossy fiber inputs^114^. Our experimental approach also does not distinguish between parvalbumin expressing axo-axonic cells and basket cells. The extent to which these PV IN subtypes and other CA3/CA2 INs such as CCK INs contribute to immature abDGC-dependent modifications of SWR properties to enhance social cognition remains to be addressed.

Taken together, we demonstrate that immature abDGCs preferentially recruit PV INs and feed-forward inhibition in DG-CA3/CA2 to decrease social memory interference and enhance social memory consolidation (**Fig. 8**). Our genetic modelling of the effective impact of neoteny of abDGCs on circuitry and cognition in adult mice suggests that protracted maturation of abDGCs expands cognitive capacity through preferential recruitment of inhibitory circuitry, a potentially testable thesis in non-human primate and human tissue^115,116^. Neoteny of abDGCs may reflect an evolutionary adaptation to offset the precipitous decline in adult hippocampal neurogenesis in higher mammals and potentially, also during natural aging^117,118^.

## Methods

### Animals

Adult male and female C57BL/6J mice (8–10 weeks old) were used as experimental subjects. They were housed in groups of three to four per cage under standard laboratory conditions. All mice were housed in a 12-hour light–dark cycle (7 a.m.–7 p.m.) in a vivarium room maintained at 22°C–24°C, with ad libitum access to food and water. Juvenile C57BL/6J mice of both sexes (25–38 days old) were used as social stimuli. All mice were group-housed, and experiments were conducted in accordance with procedures approved by the Institutional Animal Care and Use Committees at Massachusetts General Hospital and NIH guidelines (IACUC 2011N000084). Adult female mice (3–4 months old) were purchased from Jackson Laboratories for breeding. *Ascl1*-CreER^T2^ transgenic mouse line (*Ascl1^tm1.1(Cre/ERT2)Jejo^*/J, JAX Strain #:012882)^119^ was bred with *Bax^f/f^* mice^120^ to generate *Ascl1*-CreER^T2^; *Bax* ^f/f^ mice. *Bax*^f/f^ mice were generated by interbreeding *Ascl1*-CreER^T2^;*Bax*^f/f^ and *Bax*^f/f^ mice and were used to assess CreER^T2^- independent effects of tamoxifen (TAM) on behavior. Ai14 (B6;129S6-Gt(ROSA)26Sortm14(CAG-tdTomato)Hze/J) conditional reporter line was obtained from Jackson Labs, (JAX Strain #:007914)^81^ and used in all experiments where abDGCs were genetically labeled. (*Ascl1*-CreER^T2^;*Bax*^f/+^;ROSA26fSTOPtdTomato/+ mice were bred with *Bax* ^f/+^; ROSA26fSTOPtdTomato/+ mice).

### Tamoxifen injections

To induce CreER^T2^-mediated recombination of conditional alleles in this study, mice were given 3 mg of TAM intraperitoneally, once daily, for 3 consecutive days. A 10 mg/ml tamoxifen (Sigma, T-5648) solution was prepared in corn oil containing 10% ethanol. Mouse lines were obtained from Jackson Laboratories, and details are provided in Reporting Summary.

### Stereotactic viral injections

Mice received carprofen (5 mg/kg subcutaneously, Patterson Veterinary Supply) before surgery and were then anesthetized with ketamine and xylazine (10 mg/mL and 1.6 mg/mL, intraperitoneally). Mice were placed in a stereotaxic apparatus, and small holes were drilled at each injection site using the Foredom K.1070 High Speed Rotary Micromotor Kit. Bilateral injections were performed using a micro injection machine (Nanoject III, Drummond Scientific Company, PA, USA) and a micro glass pipette. The pipette was slowly lowered into the target sites and left in place for 8 minutes prior to infusion, which occurred at a rate of 2 nL/sec. The coordinates relative to bregma were: dorsal DG: 1.8 mm (AP), ±1.35 mm (ML), 2.25 mm (DV), and dorsal CA2/CA3: 1.8 mm (AP), ±2.6 mm (ML), 2.2 mm (DV). The glass pipettes were slowly withdrawn after infusion, and the skin above the incision was sutured with coated Vicryl sutures (Ethicon US LLC) and Superbond (Superbond information). Mice were monitored and received daily injections of carprofen (5 mg/kg, intraperitoneally) for 3 days following surgery.

Behavioral procedures

### Social recognition memory interference (SRMi) task

Social recognition was tested using a social discrimination procedure adapted from prior work in rats^121^ and mice ^122^. Briefly, experimental subjects were separated by transferring them to new cages (same size as their home cage) with fresh bedding for at least 2 hours before starting the session. A social discrimination session consisted of two 5-minute exposures to juveniles to the adult mouse in it’s homecage, conducted under dimmed lighting conditions (approx. 200 lx). During the "sampling" exposure, a juvenile (same sex) was introduced to the adult animal. Afterward, the juvenile was removed and kept individually in a fresh cage with food and water available ad libitum. For the "interference" session, a novel juvenile mouse was introduced to the adult 3 hours before the sampling session. After a retention interval of 24 hours, the same "sampling" juvenile was reintroduced to the adult ("choice" session), along with an additional, previously unpresented juvenile of the same strain. The duration of investigatory behavior of the adult toward each juvenile was measured separately by a trained observer blind to the animals’ treatment. A significantly longer investigation duration of the novel juvenile compared to the previously encountered one during the choice session is taken as evidence of intact recognition memory. After the end of each choice session, the experimental mice were housed in their original groups.

### Behavior procedure for DREADD virus injection group

Two weeks after viral injection surgery, mice were handled for 3 days prior to behavioral experiments to habituate them to human handling and transportation from vivarium to behavioral testing rooms. For chemogenetic manipulations of PV INs, mice were habituated to an empty syringe for 3 days and then CNO (1 mg/kg for Gi, 10mg/kg for Gq) was injected 30 min prior to experiments.

### Contextual fear conditioning

Contextual fear conditioning (CFC) occurred in conditioning chambers (18 cm x 18 cm x 30 cm; Coulbourn Instruments), each comprised of two clear Plexiglas walls and ceiling, two metal walls, a houselight, and a stainless-steel grid floor, which was connected to a shock delivery device (Coulbourn Instruments). Conditioning chambers were housed in larger, ventilated cabinets (Coulbourn Instruments). Digital cameras (Sentech) were mounted above the conditioning chambers and interfaced via USB (Actimetrics) along with the shock delivery devices to a computer running FreezeFrame software (Actimetrics) for filming, timed delivery of foot shocks, and quantifications of freezing behaviors. 8 (2M/6F) *i*Bax^Ascl1^ and 12 (8M/4F) *i*Bax^WT^ were individually handled (∼30 sec per mouse) by experimenters on three separate occasions across three consecutive days prior to undergoing CFC. Mice were transported from the vivarium and allowed to acclimate to the behavioral testing room in their homecages for at least 30 min prior to the start of any phase of CFC. At least 5 min prior to being placed in testing chamber, mice were individually housed in clean, temporary homecages before and after behavioral testing (before being returned to their group housing). Mice were randomly assigned to one of four testing chambers and were returned to the same chamber and context throughout the tests. CFC occurred in a single context (Context A). For Context A, chambers were cleaned and wiped down with 70% EtOH, the grid floor was exposed, the houselight was turned on, the fan of the cabinet was turned on, and the door of the external cabinet was closed. Mice were conditioned and tested in squads of four mice at a time (counterbalanced for group assignment, when possible), with males being tested first, then females. On day 1 of CFC, mice were placed in Context A and after a 3-min baseline, mice received three 2 sec, 0.7 mA foot shocks separated by 1-min intertrial intervals (mice remained in the chamber for 1 min after the final shock). 24 hours after the conditioning session, mice received a 5-min retrieval session in Context A. 4 weeks after a series of tamoxifen injections, mice experienced a remote retrieval session in Context A lasting 5 min. Freezing behavior (plotted in figures as a percentage of time) was automatically measured using the trial viewer function in FreezeFrame software. Specifically, and while blind to group assignments, the threshold value for each video was set to the bottom of the first trough of the motion index waveform generated by FreezeFrame (typically a value ranging from 5 to 50). Additionally, we set the minimum bout duration of freezing to be 1 sec or more.

### Immunohistochemistry

Mice were anesthetized with ketamine and xylazine (10 mg/mL and 1.6 mg/mL, IP), transcardially perfused with 4% PFA, and brains were incubated in 4% PFA at 4C overnight. Brains were placed in 30% sucrose/PBS for 2 days and then embedded in medium (OCT, Fisher HealthCare). 35 mm cryosections were obtained (Leica) and stored in PBS (0.01% sodium azide) at 4C. For immuno staining, floating sections were permeabilized, blocked in blocking solution for 2 h (PBS containing 0.3% Triton X-100 and 10% normal donkey serum, NDS), and followed by incubation with primary antibodies (PBS containing 10% NDS) at 4C overnight. Sections were then washed with PBS 3 times,10 min each, followed by incubated with secondary antibodies in PBS for 2h at room temperature (RT). Sections were then washed with PBS 3 times, 10 min each, mounted on glass slides and cover slipped with mounting medium containing DAPI. See key resources table for primary antibodies information and dilutions. For quantification of abDGCs, 6 images from 3 sections of the dorsal hippocampus were acquired with an epifluorescence microscope (Nikon) using a 10× objective. Unbiased and blinded manual quantification was used to quantify tdTomato positive abDGCs in the granule cell layer of the DG. Image analysis of PV puncta were obtained from 6 sections per mouse hippocampus. A Leica SP8 confocal laser microscope and LAS software were used to capture images in the stratum lucidum at high-resolution (2,048). For PV puncta, single confocal plane images were captured in the CA2 and CA3ab subfields using a 63X oil objective with 43X digital zoom. Quantification sample size: for PV puncta, density was averaged from 18 images per mouse. PV puncta was analyzed using the StarDist 2Dplugin and particle analysis tools in FIJI ImageJ. Threshold values were held constant across images. This protocol has been validated with electrophysiology^78^.

### Ex vivo Electrophysiology

Mice were unilaterally injected with 0.3 μL AAV-S5E2-tdTomato into dorsal CA3/CA2 and 0.2 μL AAV5-CamKII-ChR2-eYFP into the dorsal DG. 1.5 weeks after viral infusion, mice were anaesthetized with ketamine and xylazine (10 mg/ml and 1.6 mg/ml, i.p.) then transcardially perfused with ice-cold (4 °C) choline chloride-based artificial cerebrospinal fluid (ACSF) composed of (in mM): 92 choline chloride, 2.5 KCl, 1.25 NaH2PO4, 30 NaHCO3, 20 HEPES, 25 glucose, and 10 MgSO4·7H2O. Their brains were rapidly extracted following decapitation. Coronal slices (300 μm thick) containing the dorsal hippocampus were cut in ice-cold (4 °C) choline chloride ACSF using a Leica VT1000 vibratome (Leica) and transferred to warm (33 °C) normal ACSF for 30 min. Normal ACSF contained (in mM): 124 NaCl, 2.5 KCl, 1.25 NaH2PO4, 24 NaHCO3, 5 HEPES, 12.5 glucose, 2 MgSO4·7H2O, 2 CaCl2·2H2O. All ACSF solutions were adjusted to a pH of 7.4, mOsm of 305, and were saturated with carbogen (95% O2 and 5% CO2). Slices were allowed to cool to room temperature (20-22 °C) for 1 hour before recordings.

Whole-cell patch-clamp recordings were obtained using a Multiclamp 700B amplifier (Molecular Devices) low-pass filtered at 1.8 kHz with a four-pole Bessel filter and digitized with a Digidata 1550B (Molecular Devices). Slices were placed in a submersion chamber and continually perfused (>2 mL/min) with normal ACSF. Neurons were visually identified by infrared differential interference contrast imaging combined with epifluorescence using LED illumination (pE-300white, CoolLED). Pyramidal neurons in CA2 and CA3ab were distinguished by their anatomical location and distinct electrophysiological properties. Borosilicate patch pipettes had an impedance of 4-5 MΩ and filled with an internal solution containing (in mM): 120 CsMeS, 4 MgCl2, 1 EGTA, 10 HEPES, 5 QX-314, 0.4 Na3GTP, 4 MgATP, 10 phosphocreatine, 2.6 biocytin, pH 7.3, 290 mOsm. Once GΩ seal was obtained, neurons were held in voltage-clamp configuration at -70 mV and the input resistance, resting membrane potential, and capacitance were measured. Series resistance (<30 MΩ) was monitored throughout recordings and recordings were discarded if series resistance changed by >20% from baseline.

Excitatory and inhibitory postsynaptic current (EPSC and IPSC) were optically evoked with 2 ms 473 light pulses delivered above the mossy fiber pathway – the hilus of the DG. Current responses were recorded at 1.5 × threshold, defined as the minimum stimulation intensity required to produce a consistent current response beyond baseline noise. Isolation of EPSC was done by voltage clamp at -70 mV and IPSC at 0 mV. Paired pulse stimulation was evoked 5 times at in interval of 10 sec. The interevent interval between pulse stimulation was 100 ms.

Miniature EPSC and IPSC (mEPSC and mIPSC) were recorded after bath application of tetrodotoxin (TTX, 1 µM). Isolation of mEPSC was done by voltage clamp at -70 mV and mIPSC at 0 mV. Autodetection parameters for inclusion of miniature events was determined by calculating minimum threshold: Root mean square (RMS)2 × 1.5. Data acquisition was performed using Clampex and analyzed with Clampfit (Molecular Devices) and EasyElectrophysiology (V2.5.2) software.

### In vivo Electrophysiology

#### Electrode implant surgery and electrode location confirmation

Subjects were 10-11 adult male and female mice weighing approximately 30 g. Mice were anaesthetized with 1.75% isoflurane and placed in a stereotaxic frame to implant three bone screws, a recording electrode and a ground electrode. 32-site linear silicon probe array or 64ch multi shank linear silicon probe array with 30-µm diameter recording sites and 50-µm inter-site spacing (Neuronexus; part no.: A1x32-6mm-50-703 or H16x4) was placed in the dorsal hippocampus (2 mm posterior, 1 0 mm lateral; −1.8–2.3 mm ventral from Bregma) to span all layers of hippocampus. For tetrode wire recording, a four-wire stimulating electrode bundle was made by twisting together 4 × 75-µm diameter nichrome wires (California Fine Wire). The bundle was cut at an angle so as to span 0.5 mm. 4 different sites were chosen for twisted wire implant: The tips of the twisted wire were located in bilateral CA2, right side CA1 and DG. The twisted electrodes were each attached to a pin in a Mill-Max connector. After confirming that the recording electrode array extended through CA1, the wires and connectors were fixed in place with dental cement. The Omnetics connector of the recording electrode and the Mill-Max connector of the stimulating bundle were anchored to the skull along with bone screws using dental cements (C&B Metabond, Parkel and TEETs Denture Material, Cooralite Dental Mfg). A digitization board with 32 unipolar inputs (RHD2132, Intan Technologies) was connected directly to the recording electrode assembly for signal amplification and digitization. A lightweight cable (Intan Technologies) transmitted digital data to the computer using a recording system (Open Ephys, Lisbon, Portugal) that was connected to the USB port of a personal computer. The cable was connected through a lightweight swivel to enable free movement of the animal.

#### Sleep State classification

Movement was identified using accelerometer data. Epochs with accelerometer signal greater than 0.02 z were classified as awake. Stationary periods with high delta (0.5–4 Hz) power were thereafter classified as non-rapid eye movement (NREM) sleep and those with high relative theta power (a ratio of theta (5– 11 Hz) power to the sum of delta and alpha (12–30 Hz) power) as REM sleep^123^. NREM epochs < 30ms were discarded while detecting ripples as described below. When comparing sleep before and after social interaction, the last 4 NREM epochs of the preceding sleep period were compared to the first 4 NREM epochs after social interaction to ensure that the sleep periods directly preceding and following interaction were compared.

#### Detection of Ripples

Local Field Potentials (LFP) were down sampled to 2000 Hz. LFP during NREM was extracted and filtered in 100 – 200 Hz band and z-scored. Events with peak power > 4 Z were identified as potential ripple events. Thereafter events with a duration of < 30ms and inter-ripple interval < 15ms were discarded.

#### Histology

Upon conclusion of the in vivo electrophysiology experiments, all mice were transcardially perfused with 1× phosphate buffered saline (PBS) followed by 4% PFA. The brains were extracted and stored in formalin overnight. The brains were stored in 30% sucrose in 1× PBS until they were cut on a cryostat (40 µm) and thaw mounted onto gelatin-coated slides. The sections were dried for 1–2 h at room temperature and then Nissl stained. The slides were scanned at 10× with an Olympus VS120 microscope and the images were subsequently examined for electrode tracks to verify the stimulation and recording locations.

### Sex as a biological variable

We did not make any a priori predictions for sex differences. Male and female mice were included in most of the experiments in this study. Attempts were made to balance sex within each experiment and group, but we acknowledge that the ratio of males and females are not consistent across every group, and that group sizes may be underpowered for statistical comparisons of sex.

### Statistical Analysis

We adhered to accepted standards for rigorous study design and reporting to maximize the reproducibility and translational potential of our findings as described in ARRIVE Guidelines. All experimenters were blind to treatment conditions throughout data collection, scoring, and analyses. Statistical analyses were conducted using Prism v9 (GraphPad) and the minimum sample size was determined based on prior experience, existing literature, and a power analysis. Statistical significance was defined as *p*<0.05. Two tailed Student’s t tests were used for two-group comparisons. Analysis of variance (ANOVA) were used for three or more group comparisons. Repeated-measures ANOVA were used for comparison of groups across treatment condition or time. Appropriate nonparametric tests were used when datasets failed to meet parametric assumptions. Detailed statistical analyses can be found in Supplementary Table 1.

## Author contributions

A.S conceived and administered the project, A.S, A.C and J.B.A co-developed the project, A.C,L.E, T.D.G and S.M.M contributed to design, execution and analysis of behavioral paradigms, L.E and I.M performed immunohistochemistry, J.B.A performed ex vivo electrophysiology, A.C, J.B.A and L.E performed in vivo electrophysiological recordings, A.C, M.G and O.J.A performed SWR analysis, O.J.A and A.S provided supervision and funding, A.S, A.C and J.B.A wrote the manuscript with input from co-authors.

## Acknowledgements

We thank Sahay lab members, L.B.B.S and L.M.S.S for help with editing. A.S. acknowledges support from The Simons Collaboration on Plasticity and the Aging Brain, NIH R01MH111729, R01MH131652, R01MH111729-04S1, R01AG076612, R01AG076612-S1 diversity supplement, James and Audrey Foster MGH Research Scholar Award and MGH Department of Psychiatry. J.A is supported by a R01AG076612-S1 diversity supplement and The Simons Collaboration on Plasticity and the Aging Brain. T.D.G is a recipient of Brain & Behavior Research Foundation Young Investigator Award, a Harvard Brain Initiative Travel Grant, and a NIH K99/R00 Pathway to Independence Award (K99MH132768). O.J.A. acknowledges support from NIH R01MH129282, NIH R34NS127101, NIH P50NS123067, and Alzheimer’s Association Grant AARG-NTF-21-846572. This study is dedicated to CRM.

## Ethics Declarations

The authors declare no competing interests.

## Data and Code availability

All data, code and materials used in this study are available in some form to any researcher for purposes of reproducing or extending the analysis.

## Supplementary information

Extended Data Fig. 1 to 4

Supplementary Table 1

---

**Extended Data Figure 1.**
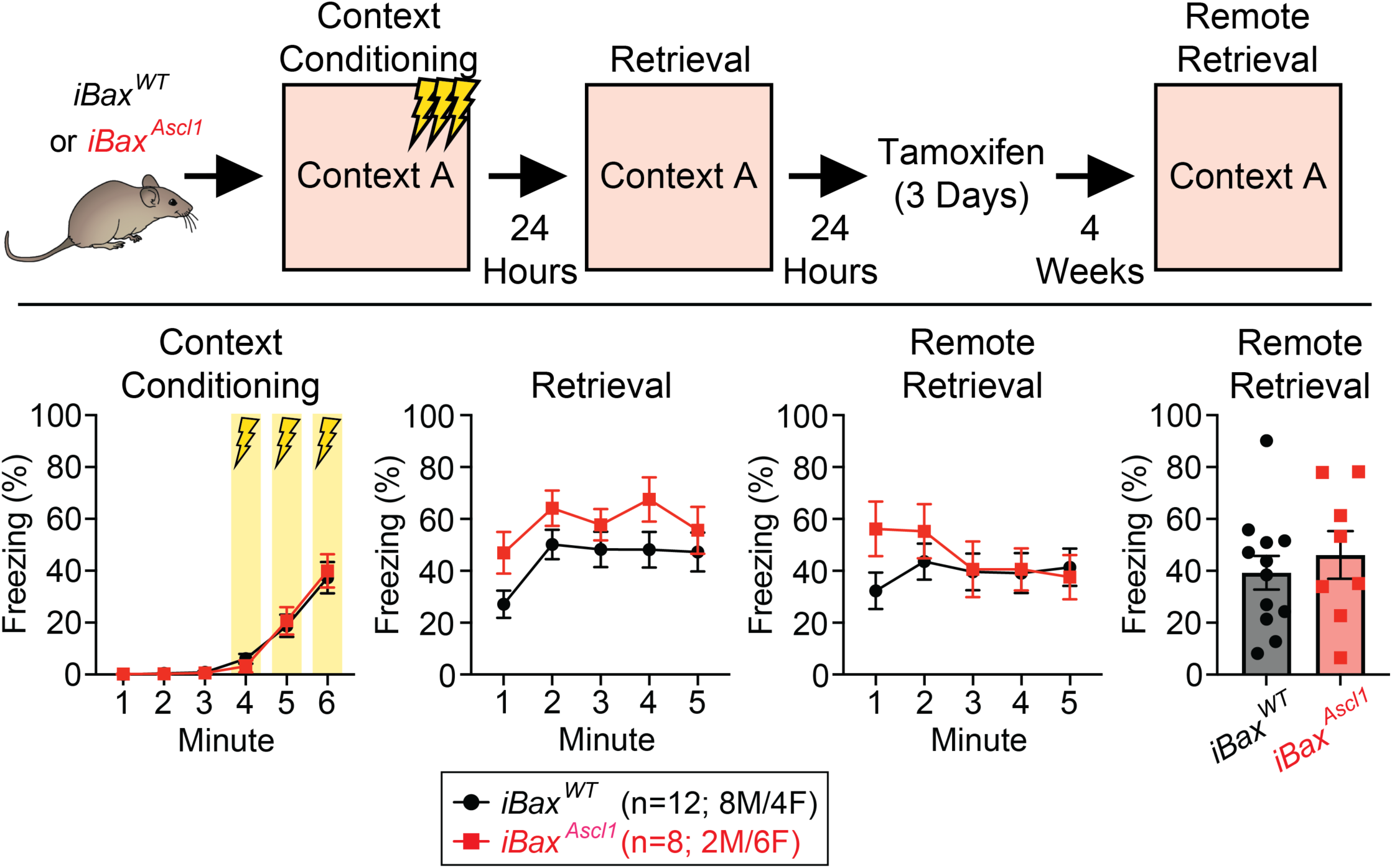
Neotenic expansion of abDGC population after learning does not promote forgetting of contextual fear memory. (Top Panel) Summary schematic for behavioral timeline. (Bottom Panels) Freezing as a percentage of time across each phase of contextual fear conditioning. There was statistically significant main effect of time in context conditioning and retrieval after 24 hours. There was significant time x group interaction in the post-induction remote retrieval, however Bonferroni’s post hoc comparisons revealed no significant differences between groups. Two-way RM ANOVA with Geisser-Greenhouse correction and Bonferroni’s posthoc comparisons were used between groups. Two-tailed unpaired *t* test was used to analyze remote retrieval summary. n = 8 *i*Bax^Ascl1^ and 12 *i*Bax^WT^ mice, significance = *p* < 0.05.

**Extended Data Figure 2.**
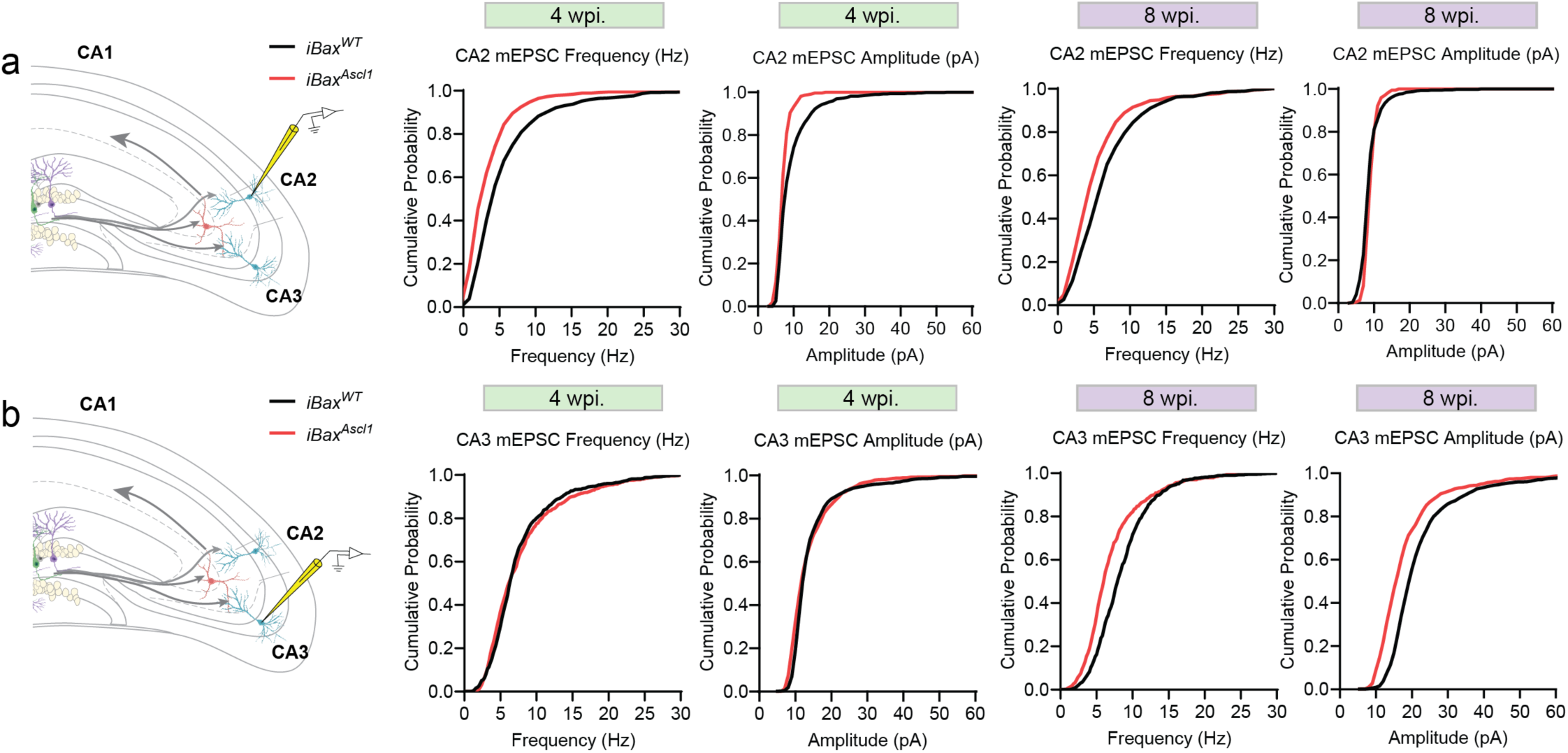
Miniature excitatory postsynaptic current from 4- and 8-week iBax^Ascl1^ mice. **a,** Schematic depicting whole-cell patch-clamp recording of mEPSC from CA2 PNs (left). Cumulative probability plots of mEPSC frequency and amplitude from PNs in 4- and 8-week post TAM injection. Kolmogorov-Smirnov test were used between groups, n = 9-13 cells, 2-4 cells per mouse, 3-4 mice per group. **b,** Schematic depicting whole-cell patch-clamp recording of mEPSC from CA3 PNs (left). Cumulative probability plots of mEPSC frequency and amplitude from PNs in 4- and 8-week post TAM injection. Kolmogorov-Smirnov test were used between groups, n = 8-14 cells, 1-4 cells per mouse, 3-4 mice per group.

**Extended Data Figure 3.**
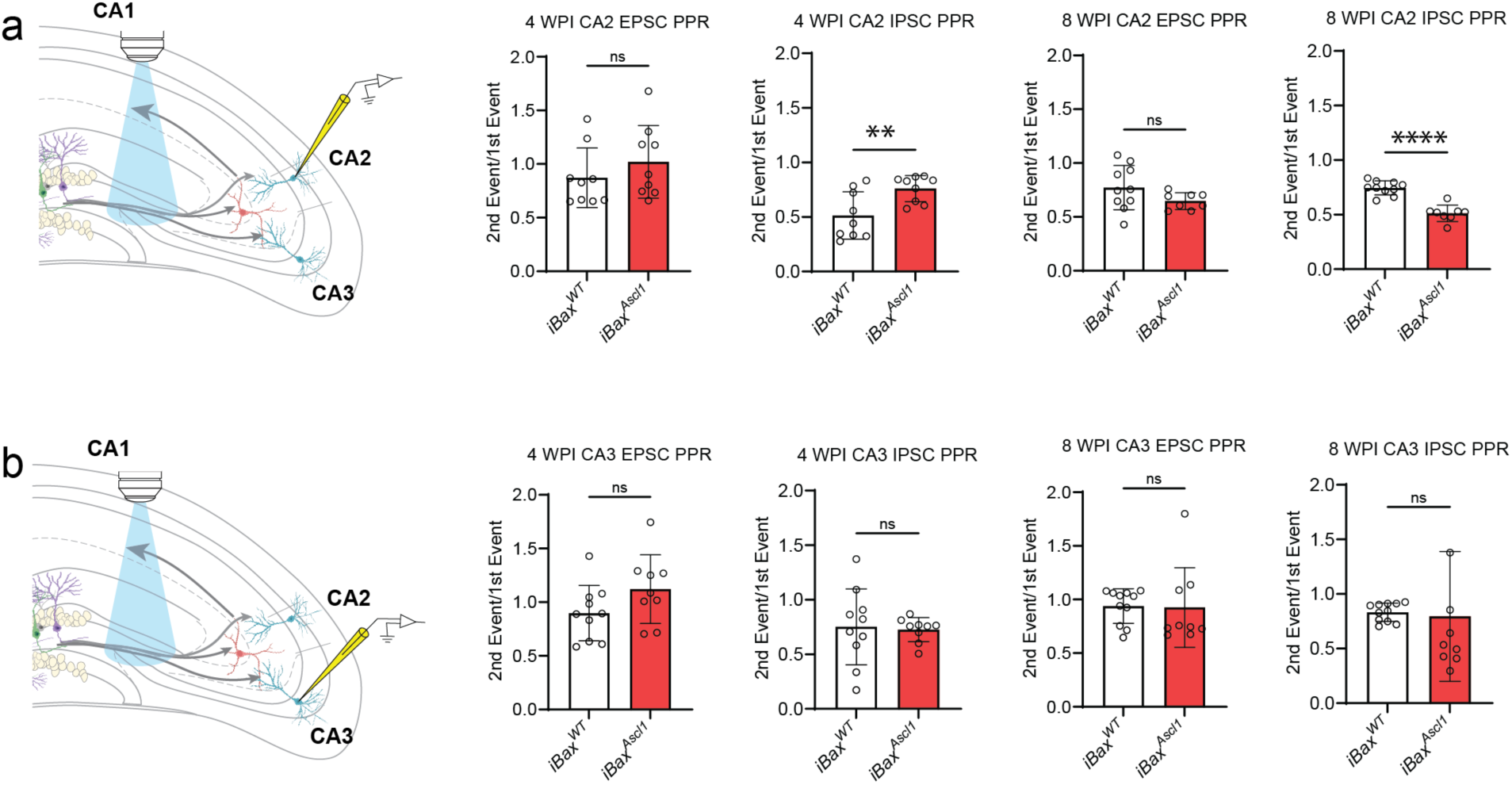
EPSC and IPSC responses to optically evoked paired pulse stimulation. **a,** Schematic depicting whole-cell patch-clamp recording of EPSC and IPSC responses to optically evoked paired pulse stimulation from CA2 PNs (left). Bar graphs depict EPSC and IPSC paired pulse ratios, the amplitude of the second event divided by the amplitude of the first event. Two-tailed unpaired *t* test was used between groups, n = 9-10 cells, 2-4 cells per mouse, 3-4 mice per group, \**p* < 0.05, Mean ± SD. **b,** Schematic depicting whole-cell patch-clamp recording of EPSC and IPSC responses to optically evoked paired pulse stimulation from CA3 PNs (left). Bar graphs depict EPSC and IPSC paired pulse ratios, the amplitude of the second event divided by the amplitude of the first event. Two-tailed unpaired *t* test was used between groups, n = 9-10 cells, 2-4 cells per mouse, 3-4 mice per group, Mean ± SD.

**Extended Data Figure 4.**
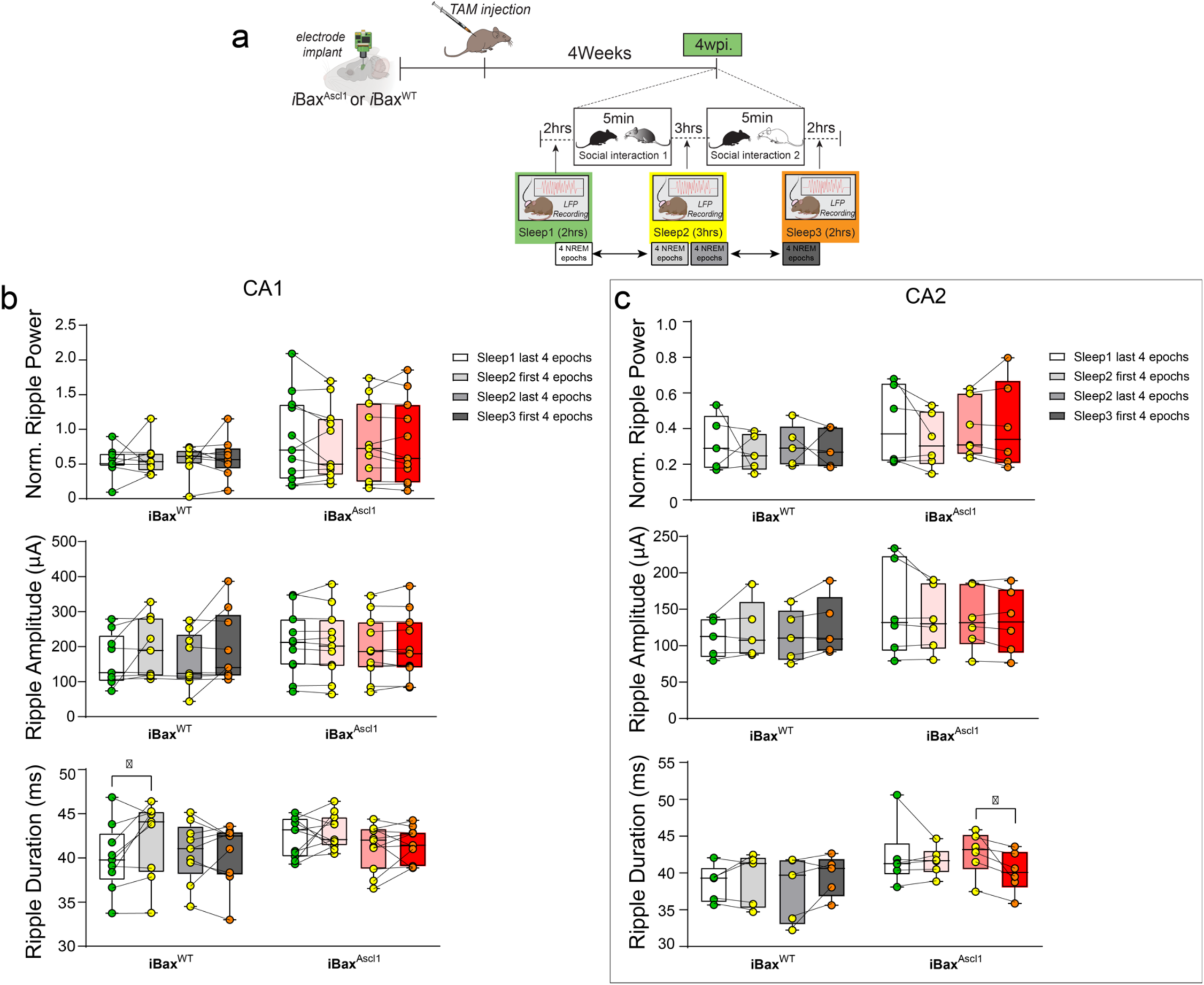
Social experience did not affect ripple power or amplitude in either group but slightly altered ripple duration. **a,** Experimental schedules: Electrodes were chronically implanted in iBax^WT^ and iBax^Ascl1^ mice. After recovery, TAM was injected into both the iBax^WT^ and iBax^Ascl1^ groups. Baseline local field potential (LFP) signals were recorded for 2 hours during sleep, 4 weeks after TAM injection (Sleep 1). Following a 5-minute social interaction, all mice underwent a 3-hour home cage sleep session (Sleep 2). This was followed by another 5-minute social interaction with a novel mouse and a subsequent 2-hour sleep session (Sleep 3). **b,** After the first 5-minute exposure to a novel mouse, neither the iBax^WT^ nor the iBax^Ascl1^ groups exhibited a significant increase in CA1 ripple power or amplitude. However, only the iBax^WT^ group showed a significant increase in ripple duration after the first exposure to the novel juvenile mouse (iBax^WT^: n = 9, iBax^Ascl1^ : n = 11). A paired t-test was used to analyze changes in ripple rate before and after social exposure. *p < 0.05. **c,** In the iBax^Ascl1^ group, CA2 ripple power, amplitude, and duration did not significantly increase after the first social interaction. However, after the second exposure to a novel mouse, CA2 ripple duration decreased in iBax^Ascl1^ (iBax^WT^: n = 5, iBaxAscl1: n = 6). A Wilcoxon matched-pairs signed-rank test was used to analyze changes in ripple rate before and after social exposure. *p < 0.05

## References

1 Altman, J. & Das, G. D. Autoradiographic and histological evidence of postnatal hippocampal neurogenesis in rats. J Comp Neurol 124, 319–335 (1965).

2 Seri, B., Garcia-Verdugo, J. M., McEwen, B. S. & Alvarez-Buylla, A. Astrocytes give rise to new neurons in the adult mammalian hippocampus. J Neurosci 21, 7153–7160 (2001).

3 van Praag, H. et al. Functional neurogenesis in the adult hippocampus. Nature 415, 1030–1034 (2002). 10.1038/4151030a 4151030a [pii]

4 Eriksson, P. S. et al. Neurogenesis in the adult human hippocampus. Nat Med 4, 1313–1317 (1998).

5 Spalding, K. L. et al. Dynamics of hippocampal neurogenesis in adult humans. Cell 153, 1219–1227 (2013). 10.1016/j.cell.2013.05.002

6 Elliott, T. et al. Hippocampal neurogenesis in adult primates: a systematic review. Mol Psychiatry (2024). 10.1038/s41380-024-02815-y

7 Toni, N. & Schinder, A. F. Maturation and Functional Integration of New Granule Cells into the Adult Hippocampus. Cold Spring Harbor perspectives in biology 8, a018903 (2015). 10.1101/cshperspect.a018903

8 Miller, S. M. & Sahay, A. Functions of adult-born neurons in hippocampal memory interference and indexing. Nat Neurosci (2019). 10.1038/s41593-019-0484-2

9 Snyder, J. S., Kee, N. & Wojtowicz, J. M. Effects of adult neurogenesis on synaptic plasticity in the rat dentate gyrus. J Neurophysiol 85, 2423–2431 (2001).

10 Schmidt-Hieber, C., Jonas, P. & Bischofberger, J. Enhanced synaptic plasticity in newly generated granule cells of the adult hippocampus. Nature 429, 184–187 (2004).

11 Zhao, C., Teng, E. M., Summers, R. G., Jr., Ming, G. L. & Gage, F. H. Distinct morphological stages of dentate granule neuron maturation in the adult mouse hippocampus. J Neurosci 26, 3–11 (2006).

12 Ge, S., Yang, C. H., Hsu, K. S., Ming, G. L. & Song, H. A critical period for enhanced synaptic plasticity in newly generated neurons of the adult brain. Neuron 54, 559–566 (2007).

13 Ge, S., Sailor, K. A., Ming, G. L. & Song, H. Synaptic integration and plasticity of new neurons in the adult hippocampus. The Journal of physiology 586, 3759–3765 (2008).

14 Toni, N. et al. Synapse formation on neurons born in the adult hippocampus. Nat Neurosci 10, 727–734 (2007).

15 Toni, N. et al. Neurons born in the adult dentate gyrus form functional synapses with target cells. Nat Neurosci 11, 901–907 (2008). https://doi.org/nn.2156 [pii] 10.1038/nn.2156

16 Sahay, A. et al. Increasing adult hippocampal neurogenesis is sufficient to improve pattern separation. Nature 472, 466–470 (2011). 10.1038/nature09817

17 Vivar, C. et al. Monosynaptic inputs to new neurons in the dentate gyrus. Nature communications 3, 1107 (2012). 10.1038/ncomms2101

18 Gu, Y. et al. Optical controlling reveals time-dependent roles for adult-born dentate granule cells. Nat Neurosci 15, 1700–1706 (2012). 10.1038/nn.3260

19 Kheirbek, M. A., Tannenholz, L. & Hen, R. NR2B-dependent plasticity of adult-born granule cells is necessary for context discrimination. J Neurosci 32, 8696–8702 (2012). 10.1523/JNEUROSCI.1692-12.2012

20 Restivo, L., Niibori, Y., Mercaldo, V., Josselyn, S. A. & Frankland, P. W. Development of Adult-Generated Cell Connectivity with Excitatory and Inhibitory Cell Populations in the Hippocampus. J Neurosci 35, 10600–10612 (2015). 10.1523/JNEUROSCI.3238-14.2015

21 Bergami, M. et al. A critical period for experience-dependent remodeling of adult-born neuron connectivity. Neuron 85, 710–717 (2015). 10.1016/j.neuron.2015.01.001

22 Goncalves, J. T. et al. In vivo imaging of dendritic pruning in dentate granule cells. Nat Neurosci 19, 788–791 (2016). 10.1038/nn.4301

23 Kennedy, W. M., Gonzalez, J. C., Lee, H., Wadiche, J. I. & Overstreet-Wadiche, L. T-Type Ca(2+) Channels Mediate a Critical Period of Plasticity in Adult-Born Granule Cells. J Neurosci 44 (2024). 10.1523/JNEUROSCI.1503-23.2024

24 Marin-Burgin, A., Mongiat, L. A., Pardi, M. B. & Schinder, A. F. Unique processing during a period of high excitation/inhibition balance in adult-born neurons. Science 335, 1238–1242 (2012). 10.1126/science.1214956

25 Ikrar, T. et al. Adult neurogenesis modifies excitability of the dentate gyrus. Frontiers in neural circuits 7, 204 (2013). 10.3389/fncir.2013.00204

26 Drew, L. J. et al. Activation of local inhibitory circuits in the dentate gyrus by adult-born neurons. Hippocampus (2015). 10.1002/hipo.22557

27 Temprana, S. G. et al. Delayed coupling to feedback inhibition during a critical period for the integration of adult-born granule cells. Neuron 85, 116–130 (2015). 10.1016/j.neuron.2014.11.023

28 McAvoy, K. M. et al. Modulating Neuronal Competition Dynamics in the Dentate Gyrus to Rejuvenate Aging Memory Circuits. Neuron 91, 1356–1373 (2016). 10.1016/j.neuron.2016.08.009

29 Luna, V. M. et al. Adult-born hippocampal neurons bidirectionally modulate entorhinal inputs into the dentate gyrus. Science 364, 578–583 (2019). 10.1126/science.aat8789

30 Adlaf, E. W. et al. Adult-born neurons modify excitatory synaptic transmission to existing neurons. eLife 6 (2017). 10.7554/eLife.19886

31 Lacefield, C. O., Itskov, V., Reardon, T., Hen, R. & Gordon, J. A. Effects of adult-generated granule cells on coordinated network activity in the dentate gyrus. Hippocampus (2010). 10.1002/hipo.20860

32 Danielson, N. B. et al. Distinct Contribution of Adult-Born Hippocampal Granule Cells to Context Encoding. Neuron 90, 101–112 (2016). 10.1016/j.neuron.2016.02.019

33 Park, E. H., Burghardt, N. S., Dvorak, D., Hen, R. & Fenton, A. A. Experience-Dependent Regulation of Dentate Gyrus Excitability by Adult-Born Granule Cells. J Neurosci 35, 11656–11666 (2015). 10.1523/JNEUROSCI.0885-15.2015

34 Lodge, M. & Bischofberger, J. Synaptic properties of newly generated granule cells support sparse coding in the adult hippocampus. Behav Brain Res 372, 112036 (2019). 10.1016/j.bbr.2019.112036

35 McHugh, S. B. et al. Adult-born dentate granule cells promote hippocampal population sparsity. Nat Neurosci 25, 1481–1491 (2022). 10.1038/s41593-022-01176-5

36 Mugnaini, M., Trinchero, M. F., Schinder, A. F., Piatti, V. C. & Kropff, E. Unique potential of immature adult-born neurons for the remodeling of CA3 spatial maps. Cell reports 42, 113086 (2023). 10.1016/j.celrep.2023.113086

37 Laham, B. J., Gore, I. R., Brown, C. J. & Gould, E. Adult-born granule cells modulate CA2 network activity during retrieval of developmental memories of the mother. eLife 12 (2024). 10.7554/eLife.90600

38 Kumar, D. et al. Sparse Activity of Hippocampal Adult-Born Neurons during REM Sleep Is Necessary for Memory Consolidation. Neuron 107, 552–565 e510 (2020). 10.1016/j.neuron.2020.05.008

39 Tuncdemir, S. N. et al. Adult-born granule cells facilitate remapping of spatial and non-spatial representations in the dentate gyrus. Neuron 111, 4024–4039 e4027 (2023). 10.1016/j.neuron.2023.09.016

40 Clelland, C. D. et al. A functional role for adult hippocampal neurogenesis in spatial pattern separation. Science 325, 210–213 (2009). 10.1126/science.1173215

41 Pan, Y. W., Chan, G. C., Kuo, C. T., Storm, D. R. & Xia, Z. Inhibition of adult neurogenesis by inducible and targeted deletion of ERK5 mitogen-activated protein kinase specifically in adult neurogenic regions impairs contextual fear extinction and remote fear memory. J Neurosci 32, 6444–6455 (2012). 10.1523/JNEUROSCI.6076-11.2012

42 Nakashiba, T. et al. Young dentate granule cells mediate pattern separation, whereas old granule cells facilitate pattern completion. Cell 149, 188–201 (2012). 10.1016/j.cell.2012.01.046

43 Wang, W. et al. Genetic activation of ERK5 MAP kinase enhances adult neurogenesis and extends hippocampus-dependent long-term memory. J Neurosci 34, 2130–2147 (2014). 10.1523/JNEUROSCI.3324-13.2014

44 Epp, J. R., Silva Mera, R., Kohler, S., Josselyn, S. A. & Frankland, P. W. Neurogenesis-mediated forgetting minimizes proactive interference. Nature communications 7, 10838 (2016). 10.1038/ncomms10838

45 Berdugo-Vega, G. et al. Increasing neurogenesis refines hippocampal activity rejuvenating navigational learning strategies and contextual memory throughout life. Nature communications 11, 135 (2020). 10.1038/s41467-019-14026-z

46 Lods, M. et al. Adult-born neurons immature during learning are necessary for remote memory reconsolidation in rats. Nature communications 12, 1778 (2021). 10.1038/s41467-021-22069-4

47. Denny, C. A., Burghardt, N. S., D.M., S., Hen, R. & Drew, M. R. 4-6 week old adult-born hippocampal neurons influence novelty-evoked exploration and contextual fear conditioning. Hippocampus In Press. (2011).

48 Kohler, S. J., Williams, N. I., Stanton, G. B., Cameron, J. L. & Greenough, W. T. Maturation time of new granule cells in the dentate gyrus of adult macaque monkeys exceeds six months. Proc Natl Acad Sci U S A 108, 10326–10331 (2011). 10.1073/pnas.1017099108

49 Ngwenya, L. B., Heyworth, N. C., Shwe, Y., Moore, T. L. & Rosene, D. L. Age-related changes in dentate gyrus cell numbers, neurogenesis, and associations with cognitive impairments in the rhesus monkey. Frontiers in systems neuroscience 9, 102 (2015). 10.3389/fnsys.2015.00102

50 Moreno-Jimenez, E. P. et al. Adult hippocampal neurogenesis is abundant in neurologically healthy subjects and drops sharply in patients with Alzheimer’s disease. Nat Med 25, 554–560 (2019). 10.1038/s41591-019-0375-9

51 Tobin, M. K. et al. Human Hippocampal Neurogenesis Persists in Aged Adults and Alzheimer’s Disease Patients. Cell Stem Cell 24, 974–982 e973 (2019). 10.1016/j.stem.2019.05.003

52 Boldrini, M. et al. Human Hippocampal Neurogenesis Persists throughout Aging. Cell Stem Cell 22, 589–599 e585 (2018). 10.1016/j.stem.2018.03.015

53 Zhou, Y. et al. Molecular landscapes of human hippocampal immature neurons across lifespan. Nature 607, 527–533 (2022). 10.1038/s41586-022-04912-w

54 Ammothumkandy, A. et al. Human adult neurogenesis loss corresponds with cognitive decline during epilepsy progression. Cell Stem Cell (2024). 10.1016/j.stem.2024.11.002

55 Tosoni, G. et al. Unique transcriptional profiles of adult human immature neurons in healthy aging, Alzheimer’s disease, and cognitive resilience. bioRxiv, 2025.2001.2008.631686 (2025). 10.1101/2025.01.08.631686

56 Snyder, J. S. Recalibrating the Relevance of Adult Neurogenesis. Trends Neurosci 42, 164–178 (2019). 10.1016/j.tins.2018.12.001

57 Baker, S. et al. The Human Dentate Gyrus Plays a Necessary Role in Discriminating New Memories. Curr Biol 26, 2629–2634 (2016). 10.1016/j.cub.2016.07.081

58 Leal, S. L. & Yassa, M. A. Integrating new findings and examining clinical applications of pattern separation. Nat Neurosci 21, 163–173 (2018). 10.1038/s41593-017-0065-1

59 Berron, D. et al. Strong Evidence for Pattern Separation in Human Dentate Gyrus. J Neurosci 36, 7569–7579 (2016). 10.1523/JNEUROSCI.0518-16.2016

60 Sakon, J. J. & Suzuki, W. A. A neural signature of pattern separation in the monkey hippocampus. Proc Natl Acad Sci U S A 116, 9634–9643 (2019). 10.1073/pnas.1900804116

61 McHugh, T. J. et al. Dentate Gyrus NMDA Receptors Mediate Rapid Pattern Separation in the Hippocampal Network. Science 317, 94–99 (2007).

62 Raam, T., McAvoy, K. M., Besnard, A., Veenema, A. H. & Sahay, A. Hippocampal oxytocin receptors are necessary for discrimination of social stimuli. Nature communications 8, 2001 (2017). 10.1038/s41467-017-02173-0

63 GoodSmith, D. et al. Spatial Representations of Granule Cells and Mossy Cells of the Dentate Gyrus. Neuron 93, 677–690 e675 (2017). 10.1016/j.neuron.2016.12.026

64 Woods, N. I. et al. The Dentate Gyrus Classifies Cortical Representations of Learned Stimuli. Neuron 107, 173–184 e176 (2020). 10.1016/j.neuron.2020.04.002

65 Senzai, Y. & Buzsaki, G. Physiological Properties and Behavioral Correlates of Hippocampal Granule Cells and Mossy Cells. Neuron 93, 691–704 e695 (2017). 10.1016/j.neuron.2016.12.011

66 Neunuebel, J. P. & Knierim, J. J. CA3 retrieves coherent representations from degraded input: direct evidence for CA3 pattern completion and dentate gyrus pattern separation. Neuron 81, 416–427 (2014). 10.1016/j.neuron.2013.11.017

67 Leutgeb, J. K., Leutgeb, S., Moser, M. B. & Moser, E. I. Pattern separation in the dentate gyrus and CA3 of the hippocampus. Science 315, 961–966 (2007).

68 Gilbert, P. E., Kesner, R. P. & Lee, I. Dissociating hippocampal subregions: double dissociation between dentate gyrus and CA1. Hippocampus 11, 626–636 (2001).

69 Hainmueller, T. & Bartos, M. Dentate gyrus circuits for encoding, retrieval and discrimination of episodic memories. Nat Rev Neurosci 21, 153–168 (2020). 10.1038/s41583-019-0260-z

70 Cayco-Gajic, N. A. & Silver, R. A. Re-evaluating Circuit Mechanisms Underlying Pattern Separation. Neuron 101, 584–602 (2019). 10.1016/j.neuron.2019.01.044

71 Burghardt, N. S., Park, E. H., Hen, R. & Fenton, A. A. Adult-born hippocampal neurons promote cognitive flexibility in mice. Hippocampus 22, 1795–1808 (2012). 10.1002/hipo.22013

72 Zhang, J. et al. Ezh2 regulates adult hippocampal neurogenesis and memory. J Neurosci 34, 5184–5199 (2014). 10.1523/JNEUROSCI.4129-13.2014

73 Zhuo, J. M. et al. Young adult born neurons enhance hippocampal dependent performance via influences on bilateral networks. eLife 5 (2016). 10.7554/eLife.22429

74 Gustus, K. C. et al. Genetic inactivation of synaptosomal-associated protein 25 (SNAP-25) in adult hippocampal neural progenitors impairs pattern discrimination learning but not survival or structural maturation of newborn dentate granule cells. Hippocampus 28, 735–744 (2018). 10.1002/hipo.23008

75 Frechou, M. A. et al. Adult neurogenesis improves spatial information encoding in the mouse hippocampus. Nature communications 15, 6410 (2024). 10.1038/s41467-024-50699-x

76 Niibori, Y. et al. Suppression of adult neurogenesis impairs population coding of similar contexts in hippocampal CA3 region. Nature communications 3, 1253 (2012). 10.1038/ncomms2261

77 Tronel, S. et al. Adult-born neurons are necessary for extended contextual discrimination. Hippocampus 22, 292–298 (2012). 10.1002/hipo.20895

78 Shih, Y. T., Alipio, J. B. & Sahay, A. An inhibitory circuit-based enhancer of DYRK1A function reverses Dyrk1a-associated impairment in social recognition. Neuron 111, 3084–3101 e3085 (2023). 10.1016/j.neuron.2023.09.009

79 Shih, Y. T. et al. Pro-cognitive restoration of experience-dependent parvalbumin inhibitory neuron plasticity in neurodevelopmental disorders. Res Sq (2025). 10.21203/rs.3.rs-5624085/v1

80 Yang, S. M., Alvarez, D. D. & Schinder, A. F. Reliable Genetic Labeling of Adult-Born Dentate Granule Cells Using Ascl1 CreERT2 and Glast CreERT2 Murine Lines. J Neurosci 35, 15379–15390 (2015). 10.1523/JNEUROSCI.2345-15.2015

81 Madisen, L. et al. A robust and high-throughput Cre reporting and characterization system for the whole mouse brain. Nat Neurosci 13, 133–140 (2010). 10.1038/nn.2467

82 Perna, J. C., Wotjak, C. T., Stork, O. & Engelmann, M. Timing of presentation and nature of stimuli determine retroactive interference with social recognition memory in mice. Physiol Behav 143, 10–14 (2015). 10.1016/j.physbeh.2015.02.029

83 Gao, A. et al. Elevation of Hippocampal Neurogenesis Induces a Temporally Graded Pattern of Forgetting of Contextual Fear Memories. J Neurosci 38, 3190–3198 (2018). 10.1523/JNEUROSCI.3126-17.2018

84 Hitti, F. L. & Siegelbaum, S. A. The hippocampal CA2 region is essential for social memory. Nature 508, 88–92 (2014). 10.1038/nature13028

85 Smith, A. S., Williams Avram, S. K., Cymerblit-Sabba, A., Song, J. & Young, W. S. Targeted activation of the hippocampal CA2 area strongly enhances social memory. Mol Psychiatry 21, 1137–1144 (2016). 10.1038/mp.2015.189

86 Chiang, M. C., Huang, A. J. Y., Wintzer, M. E., Ohshima, T. & McHugh, T. J. A role for CA3 in social recognition memory. Behav Brain Res 354, 22–30 (2018). 10.1016/j.bbr.2018.01.019

87 Oliva, A., Fernandez-Ruiz, A., Leroy, F. & Siegelbaum, S. A. Hippocampal CA2 sharp-wave ripples reactivate and promote social memory. Nature 587, 264–269 (2020). 10.1038/s41586-020-2758-y

88 Chen, S. et al. A hypothalamic novelty signal modulates hippocampal memory. Nature 586, 270–274 (2020). 10.1038/s41586-020-2771-1

89 Alexander, G. M. et al. CA2 neuronal activity controls hippocampal low gamma and ripple oscillations. eLife 7 (2018). 10.7554/eLife.38052

90 Alexander, G. M. et al. Social and novel contexts modify hippocampal CA2 representations of space. Nature communications 7, 10300 (2016). 10.1038/ncomms10300

91 Alexander, G. M. et al. Remote control of neuronal activity in transgenic mice expressing evolved G protein-coupled receptors. Neuron 63, 27–39 (2009). 10.1016/j.neuron.2009.06.014

92 Vormstein-Schneider, D. et al. Viral manipulation of functionally distinct interneurons in mice, non-human primates and humans. Nat Neurosci 23, 1629–1636 (2020). 10.1038/s41593-020-0692-9

93 Dimidschstein, J. et al. A viral strategy for targeting and manipulating interneurons across vertebrate species. Nat Neurosci 19, 1743–1749 (2016). 10.1038/nn.4430

94 Klausberger, T. et al. Brain-state- and cell-type-specific firing of hippocampal interneurons in vivo. Nature 421, 844–848 (2003). 10.1038/nature01374

95 Hu, H., Gan, J. & Jonas, P. Interneurons. Fast-spiking, parvalbumin(+) GABAergic interneurons: from cellular design to microcircuit function. Science 345, 1255263 (2014). 10.1126/science.1255263

96 Schlingloff, D., Kali, S., Freund, T. F., Hajos, N. & Gulyas, A. I. Mechanisms of sharp wave initiation and ripple generation. J Neurosci 34, 11385–11398 (2014). 10.1523/JNEUROSCI.0867-14.2014

97 Vancura, B., Geiller, T., Grosmark, A., Zhao, V. & Losonczy, A. Inhibitory control of sharp-wave ripple duration during learning in hippocampal recurrent networks. Nat Neurosci 26, 788–797 (2023). 10.1038/s41593-023-01306-7

98 Girardeau, G., Benchenane, K., Wiener, S. I., Buzsaki, G. & Zugaro, M. B. Selective suppression of hippocampal ripples impairs spatial memory. Nat Neurosci 12, 1222–1223 (2009). 10.1038/nn.2384

99 Nakashiba, T., Buhl, D. L., McHugh, T. J. & Tonegawa, S. Hippocampal CA3 output is crucial for ripple-associated reactivation and consolidation of memory. Neuron 62, 781–787 (2009). 10.1016/j.neuron.2009.05.013

100 Joo, H. R. & Frank, L. M. The hippocampal sharp wave-ripple in memory retrieval for immediate use and consolidation. Nat Rev Neurosci 19, 744–757 (2018). 10.1038/s41583-018-0077-1

101 Fernandez-Ruiz, A. et al. Long-duration hippocampal sharp wave ripples improve memory. Science 364, 1082–1086 (2019). 10.1126/science.aax0758

102 Norman, Y. et al. Hippocampal sharp-wave ripples linked to visual episodic recollection in humans. Science 365 (2019). 10.1126/science.aax1030

103 Twarkowski, H., Steininger, V., Kim, M. J. & Sahay, A. A dentate gyrus-CA3 inhibitory circuit promotes evolution of hippocampal-cortical ensembles during memory consolidation. eLife 11 (2022). 10.7554/eLife.70586

104 Dranovsky, A. et al. Experience dictates stem cell fate in the adult hippocampus. Neuron 70, 908–923 (2011). 10.1016/j.neuron.2011.05.022

105 Acsady, L., Kamondi, A., Sik, A., Freund, T. & Buzsaki, G. GABAergic cells are the major postsynaptic targets of mossy fibers in the rat hippocampus. J Neurosci 18, 3386–3403 (1998).

106 Toth, K., Suares, G., Lawrence, J. J., Philips-Tansey, E. & McBain, C. J. Differential mechanisms of transmission at three types of mossy fiber synapse. J Neurosci 20, 8279–8289 (2000).

107 Ruediger, S. et al. Learning-related feedforward inhibitory connectivity growth required for memory precision. Nature 473, 514–518 (2011). 10.1038/nature09946

108 Pelkey, K. A. et al. Hippocampal GABAergic Inhibitory Interneurons. Physiol Rev 97, 1619–1747 (2017). 10.1152/physrev.00007.2017

109 Guo, N. et al. Dentate granule cell recruitment of feedforward inhibition governs engram maintenance and remote memory generalization. Nat Med 24, 438–449 (2018). 10.1038/nm.4491

110 Sahay, A., Wilson, D. A. & Hen, R. Pattern separation: a common function for new neurons in hippocampus and olfactory bulb. Neuron 70, 582–588 (2011). 10.1016/j.neuron.2011.05.012

111 Pouille, F. & Scanziani, M. Enforcement of temporal fidelity in pyramidal cells by somatic feed-forward inhibition. Science 293, 1159–1163 (2001). 10.1126/science.1060342

112 Pouille, F., Marin-Burgin, A., Adesnik, H., Atallah, B. V. & Scanziani, M. Input normalization by global feedforward inhibition expands cortical dynamic range. Nat Neurosci 12, 1577–1585 (2009). 10.1038/nn.2441

113 Akers, K. G. et al. Hippocampal neurogenesis regulates forgetting during adulthood and infancy. Science 344, 598–602 (2014). 10.1126/science.1248903

114 Hunt, D. L., Linaro, D., Si, B., Romani, S. & Spruston, N. A novel pyramidal cell type promotes sharp-wave synchronization in the hippocampus. Nat Neurosci 21, 985–995 (2018). 10.1038/s41593-018-0172-7

115 Pelkey, K. A. et al. Evolutionary conservation of hippocampal mossy fiber synapse properties. Neuron 111, 3802–3818 e3805 (2023). 10.1016/j.neuron.2023.09.005

116 Watson, J. F. et al. Human hippocampal CA3 uses specific functional connectivity rules for efficient associative memory. Cell (2024). 10.1016/j.cell.2024.11.022

117 Morgenstern, N. A., Lombardi, G. & Schinder, A. F. Newborn granule cells in the ageing dentate gyrus. The Journal of physiology 586, 3751–3757 (2008). https://doi.org/jphysiol.2008.154807 [pii] 10.1113/jphysiol.2008.154807

118 Trinchero, M. F. et al. High Plasticity of New Granule Cells in the Aging Hippocampus. Cell reports 21, 1129–1139 (2017). 10.1016/j.celrep.2017.09.064

119 Kim, E. J., Ables, J. L., Dickel, L. K., Eisch, A. J. & Johnson, J. E. Ascl1 (Mash1) defines cells with long-term neurogenic potential in subgranular and subventricular zones in adult mouse brain. PLoS ONE 6, e18472 (2011). 10.1371/journal.pone.0018472

120 Takeuchi, O. et al. Essential role of BAX,BAK in B cell homeostasis and prevention of autoimmune disease. Proc Natl Acad Sci U S A 102, 11272–11277 (2005).

121 Engelmann, M., Wotjak, C. T. & Landgraf, R. Social discrimination procedure: an alternative method to investigate juvenile recognition abilities in rats. Physiol Behav 58, 315–321 (1995). 10.1016/0031-9384(95)00053-l

122 Richter, K., Wolf, G. & Engelmann, M. Social recognition memory requires two stages of protein synthesis in mice. *Learning & memory (Cold Spring Harbor*, N.Y 12, 407–413 (2005). 10.1101/lm.97505

123 Ghosh, M. et al. Running speed and REM sleep control two distinct modes of rapid interhemispheric communication. Cell reports 40, 111028 (2022). 10.1016/j.celrep.2022.111028

